# Identification of mammalian transcription factors that bind to inaccessible chromatin

**DOI:** 10.1101/2023.03.15.532796

**Authors:** Romana T. Pop, Alessandra Pisante, Dorka Nagy, Patrick C. N. Martin, Liudmila A Mikheeva, Ateequllah Hayat, Gabriella Ficz, Nicolae Radu Zabet

**Author notes:** The authors wish it to be known that, in their opinion, the first three authors should be regarded as joint First Authors.

## Abstract

Transcription factors (TFs) are proteins that affect gene expression by binding to regulatory regions of DNA in a sequence specific manner. The binding of TFs to DNA is controlled by many factors, including the DNA sequence, concentration of TF, chromatin accessibility and co-factors. Here, we systematically investigated the binding mechanism of hundreds of TFs by analysing ChIP-seq data with our explainable statistical model, ChIPanalyser. This tool uses as inputs the DNA sequence binding motif; the capacity to distinguish between strong and weak binding sites; the concentration of TF; and chromatin accessibility. We asked whether TFs preferred to bind to DNA in open or dense chromatin conformation and found that approximately one third of TFs are predicted to bind the genome in a DNA accessibility independent fashion. Our model predicted this to be the case when the TF binds to its strongest binding regions in the genome, and only a small number of TFs have the capacity to bind dense chromatin at their weakest binding regions, such as CTCF USF2 and CEBPB. Our study demonstrated that the binding of hundreds of human and mouse TFs is predicted by ChIPanalyser with high accuracy and showed that many TFs can bind dense chromatin.

## Introduction

Site specific transcription factors (TFs) control gene expression by binding to gene promoters and enhancers (1, 2) but prediction of their binding in different cell types has been a significant challenge to date. Therefore, we do not have a clear understanding of the underlying mechanism that underpins TF binding in various biological contexts. The advent of high throughput technologies, such as chromatin immunoprecipitation followed by sequencing (ChIP-seq) drove huge progress in generating empirical data on TF binding profiles, which is now the gold standard method to experimentally determine TF binding profiles (3). Nevertheless, despite their notable impact on profiling TF binding, ChIP methods do not provide a mechanistic understanding of the binding events of TFs to the genome.

TFs bind to the DNA at specific short sequences known as *motifs*, where TFs recognise and bind their motifs with much higher affinity than any other sequence (4–6). There is a wide array of methods, both *in vitro* and *in vivo*, that can be used to determine TF binding sites (3, 7, 8). However, the presence of a motif is not sufficient for a TF to bind or regulate a gene (9–11). Thus, other factors besides DNA sequence must influence TF binding, for example TF concentration (12–15).

Furthermore, TFs in humans and other higher eukaryotes also face the challenge posed by the complex structure of chromatin. Indeed, nucleosomes have long been known to impede TF binding and nucleosome-rich regions are associated with transcriptionally inactive chromatin (16, 17). Thus, nucleosome positioning and DNA accessibility is another significant factor influencing TF binding.

While it is assumed that in general TFs cannot bind nucleosome-rich chromatin, a subset of TFs known as pioneer factors can interact with nucleosomes and bind their cognate DNA. Not only that, but they are also able to displace nucleosomes and make way for other TFs to bind without the use of ATP-dependent chromatin re-modelers (18). The first pioneer factors discovered were FOXA1 and GATA4, two TFs that play an important role in endoderm formation during embryogenesis (19, 20). Since then, several other TFs involved in a variety of processes have been identified as having pioneer function (21). Other examples are the pluripotency factors (SOX2, OCT3/4, KLF4 and c-MYC), which can reprogram somatic cells and revert them to a pluripotent state. These factors have been shown to have pioneer properties, except for c-MYC which appears to lack pioneer properties itself, but is a co-factor that enhances the activity of SOX2/OCT4 (22, 23).

Bookmarking of binding sites by TFs (i.e. the continued occupancy of TFs at their binding sites even during transcriptionally inactive phases, such as during mitosis) is a mechanism for quick reactivation of transcription sites after cell division. This behaviour has been observed in several TFs such as FOXA1 (24), GATA1 (25), as well as the pluripotency factors (26) and it serves to maintain cellular differentiation, or lack thereof, in the case of the pluripotency factors.

The mechanisms by which pioneer factors open chromatin are not well understood. While it has been shown that FOXA1-mediated chromatin relaxation does not require ATP-dependent chromatin remodelers, it is unclear whether re-modelers are recruited (18, 27). In the case of FOXA1, there is evidence for it directly causing chromatin relaxation through the displacement of linker histones (28). Other factors, such as OCT4, are known to recruit chromatin re-modelers to their binding site to facilitate chromatin opening. For example, OCT4 recruits the SWI/SNF complex of chromatin remodelers, particularly Brg1 (29–31).

When a TF binds to dense chromatin, it can either open the chromatin, stay idle or help maintain it in a closed state. If binding results in opened chromatin, the TF would be classified as a pioneer factor, while in the latter case, it can be classified either as an insulator or a bookmarking TF. Insulators can bind at the boundary between dense and open chromatin and stop the spreading of heterochromatin, thereby preventing gene silencing and gene inactivation (32). In addition, insulators can bind between the enhancer and the promoter, thus blocking their interaction and interfering with gene expression (33). One interesting example of an insulator is CTCF, which appears to have the ability to displace nucleosomes after cell division and maintain nucleosome depleted regions in some contexts (34), while other times its binding is inhibited by the presence of nucleosomes (35, 36). This indicates that the relationship between TFs and chromatin is complex and context dependent.

Here, we investigate the binding profiles and chromatin accessibility preferences of human and mouse TFs using ChIPanalyser, an explainable statistical model we previously developed (13, 37). Other tools to predict ChIP profiles (38–40) train an opaque/black box machine learning (ML) model on ChIP data and predict binding profiles in other cell types. While those models can be interpreted, it is often challenging to interpret mechanistically what drives the binding of a TF. ChIPanalyser uses a bottom-up approach, where the models start from known biological and physical components of TF binding and we train the parameters of the model on ChIP data (11, 13, 37, 41). One assumption we make is that the DNA binding motif (in the form of PWM) is an accurate representation of the binding preference of a TF. This is often the case (42), but there are also exceptions, and the ML models predict different binding motifs (38–40).

We first train the ChIPanalyser model on bulk ChIP-seq and DNA accessibility data (e.g., ATAC-seq or DNaseI-seq) data in order to estimate TF binding parameters, and then use those to determine whether TFs prefer to bind open or nucleosome associated DNA. Our results show that many TFs bind to DNA in an accessibility independent manner at their strongest binding sites. We highlight different behavioural classes among mouse and human TFs. Additionally, the ability to bind to inaccessible DNA can be linked to various functional roles such as bookmarking, chromatin opening or chromatin insulation.

## Materials and Methods

### Datasets

#### K562 cell line

To investigate TF binding behaviour, we considered 244 TFs in K562 cells for which ChIP-seq data were available from ENCODE (43, 44). First, we selected ChIP-seq datasets in WT K562 cells and excluded from further analysis data where the cells were subjected to different treatments (e.g., RNAi). Furthermore, TFs for which a PWM motif was not available in the JASPAR database and had less than 60 ChIP-seq peaks were also excluded from the dataset. All metadata information for the downloaded data can be found in supplementary data Table S1. The final number of TFs after triage was 110 (see Table S2). Where multiple experiments were available for one TF, the data were merged.

In addition to the TF ChIP-seq data, DNase I hypersensitivity data were also downloaded from ENCODE for the K562 cell line (experiment accession ENCSR000EOT). This was processed in the same way as the ChIP-seq data (see below) until the peak calling stage, where broad peaks were called with a q-value threshold of 0.1 instead.

#### Mouse cell lines

We considered 78 ChIP-seq datasets in 8 mouse cell lines from ENCODE (43, 44) and, following the same filtering steps as in the K562 cells (motif in JASPAR core and more than 60 ChIP-seq peaks), 60 ChIP-seq datasets were selected (Table S6). We used several datasets for DNA accessibility (43, 45–47)(45–48) (see Table S7).

#### IMR90 and HepG2 cell lines

We considered 11 TFs in IMR90 and 3 in HepG2 cell lines using ChIP-seq data available from ENCODE (43, 44) (see Table S9). Raw human chromatin accessibility fastq datasets for IMR90 and HepG2 (in the form of DNase-seq, MNase-seq, ATAC-seq and NOMe-seq) were downloaded from ENCODE release 3 (44).

Note that whilst ATAC-seq, DNase-seq and MNase-seq follow the same accessibility pattern (starting at 100% accessible DNA for QDA=0 and then gradually decreasing their percentage of accessible genome), NOMe-seq does not follow this pattern and in this case QDA=0 corresponds to 41% of accessible regions and QDA=0.99 corresponds to 8% accessible regions (Figure S10C-D). In other words, there are no reads from 59% of the genome in the NOMe-seq data from IMR90. In the case of IMR90 cells, MNase-seq does not reach 0%, which can be explained by the quality of the data and the size of the sequencing library (MNase-seq measures the nucleosome profiles and, thus, increasing the sequencing depth could allow capturing regions with higher density of nucleosomes and lower accessibility).

To select regions with strong, medium and weak ChIP signal, we first ordered 50 Kb regions based on the number of ChIP peaks they contain for the corresponding TF and then we selected groups of 50 regions from the top, middle and bottom of the list.

When analysing the different methods to estimate DNA accessibility, we found that ATAC-seq and DNaseI-seq showed better performance compared to MNase-seq and NOMe-seq. Since this was performed in two different cell lines with different DNA accessibility datasets, our results indicate that this would not be affected by the dataset quality. However, given the lower number of analysed TFs for the comparison of the different methods to measure accessibility, these results could also be impacted by a single lower quality dataset.

### Pre-processing of ChIP-seq and DNA accessibility data

We used Trimmomatic v. 0.39 (49) or cutadapt version 1.18 (50) to remove ILLUMINA adapters and poor-quality reads. Then, the data were aligned to the hg38 reference genome (51) with bowtie2 version 2.3.4.1 (52). Peaks were called with macs2 version 2.1.2 with a q-value threshold of 0.5 (53).

### Modelling TF binding with ChIPanalyser

#### Model description

ChIPanalyser implements an approximation of the statistical thermodynamics model and is described in detail in (54, 55). Briefly, ChIPanalyser estimates ChIP-seq like profiles based on four parameters: *(i)* a weighted DNA binding motif referred to as a position weight matrix (PWM), *(ii)* DNA accessibility data, *(iii)* the number of molecules bound to the DNA (determined experimentally or predicted) and *(iv)* a factor that modulates TF specificity. The model outputs the probability that a TF is bound to a site *j* as given by:

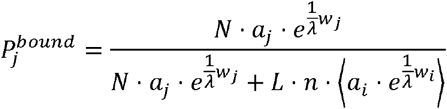

Where *N* is the number of TF molecules bound to the genome, *a_j_* is a measure of DNA accessibility at site *j* on the genome (the probability that site *j* is in accessible chromatin), *λ* is the specificity scaling factor, *w* is the PWM score and *L* and *n* are the length and ploidy of the genome, respectively.

#### Computing optimal parameters

The number of bound molecules *N* and the TF specificity factor *λ* are two of the parameters that are needed for estimating ChIP-like profiles. These are difficult to measure experimentally, so we estimated them by training ChIPanalyser on the corresponding ChIP-seq data. We binned the genome into 50kb bins and trained the model on the top 10 bins with the highest ChIP-seq signal. We then validated the estimated parameters on a different set of 50 bins of 50kb each from the same ChIP-seq dataset.

#### Quantile density accessibility (QDA)

To assess to role of chromatin accessibility in the binding of TFs, we calculated quantiles between 0-0.99 for the accessibility data and subset it based on these (55). We termed this analysis quantile density accessibility (QDA). Each QDA represents a subset of the genomic regions that the model considers accessible. For each quantile, we selected the regions with accessibility scores equal or greater than the quantile value such that a QDA of 0 corresponds to the subset of regions greater or equal to the “0 quantile” of the distribution of scores (i.e., all regions), a QDA of 0.5 corresponds to the subset of regions with scores greater or equal to the median of the distribution (top 50% regions with highest DNA accessibility signal) and a QDA of 0.9 corresponds to the subset of regions with scores greater than the 90^th^ percentile of the distribution (top 10% regions with highest DNA accessibility signal) (Figure 1B-C).

**Figure 1.**
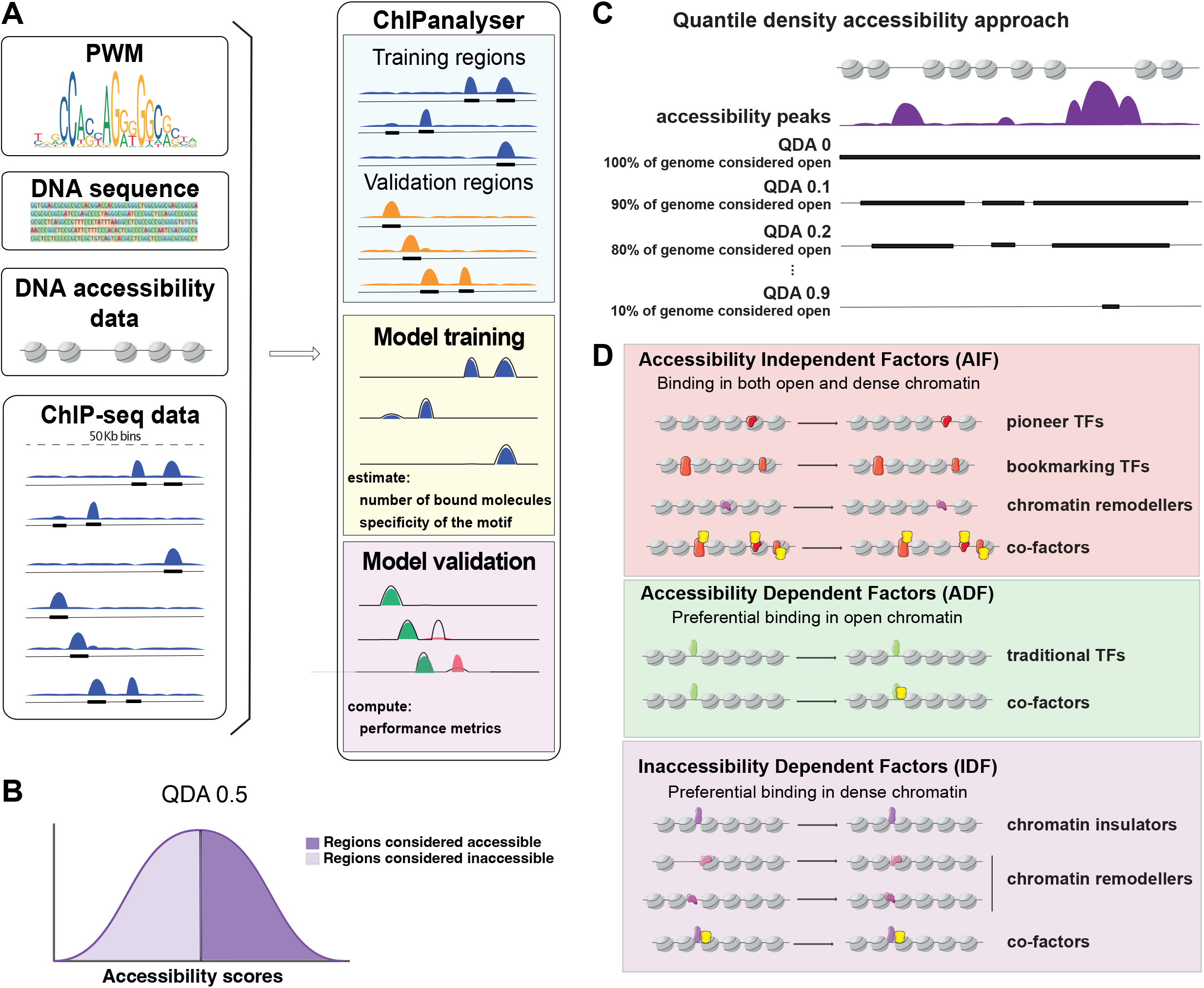
Workflow overview. (A) ChIPanalyser models TF binding based on PWM motifs, DNA sequence, and DNA accessibility data together with ChIP-seq data. After extracting TF binding profiles from ChIP-seq data, we split the genomic regions into 50 Kb bins and selected the top 10 regions with the highest number of ChIP-seq peaks as training regions, and the following 50 regions with the highest number of ChIP-seq peaks as validation regions. The ChIPanalyser model was trained on the training regions to minimise mean squared error (MSE), then validated to estimate the accuracy of the binding profile predictions. (B-C) Approach to investigate TF preference for DNA accessibility. (B) DNA accessibility data (DNaseI-seq or ATAC-seq) is used to (C) select the regions of the genome that have a signal above a threshold (e.g., QDA of 0.1 results in selecting the 90% of the genome with the highest levels of accessibility signal). (D) Graphical representation of the binding properties of each class of TF and how each class affects chromatin structure after binding. Accessibility Independent Factors (AIFs) bind open or dense chromatin without preference, but AIF binding sometimes can displace nucleosomes. Accessibility Dependent Factors (ADFs) bind open chromatin only and no changes in accessibility are observed after binding. Finally, Inaccessibility Dependent Factors (IDFs) bind chromatin in three different scenarios:*(i)* bind dense chromatin and maintain it (no changes are observed); *(ii)* bind dense chromatin and reinforce it, thus changes are observed in the increased number of nucleosomes; and lastly *(iii)* bind open chromatin, which becomes compacted. The different classes of TFs that group as AIFs (pioneer, bookmarking, chromatin remodeller and co-factors of those), ADFs (traditional TFs and their co-factors) and IDFs (insulators, chromatin remodeller and co-factors of those) are shown in the side panels.

### Clustering of the TFs

To cluster TFs based on their preference for open or dense chromatin, we first used k-means clustering to identify the different classes of TFs from the data without prior assumptions. Our analysis showed that there are two main classes of TFs (Figure S3A), namely: *(i)* TFs displaying accessibility independent binding (62) and *(ii)* TFs displaying a clear accessibility dependent binding (48). We did not find TFs that display inaccessibility dependent binding, which indicates that either there are fewer TFs displaying this behaviour, or that our collection of ChIP-seq data does not contain many TFs displaying that behaviour. Nevertheless, four TFs showed higher accuracy of predictions when assuming they can bind anywhere in the genome, compared when their binding is restricted only to accessible regions (see purple TFs in Figure S3B-C). This indicates that some of the classes in Figure 1B have too few TFs in our dataset to be detected by k-means clustering.

Since only a few TFs displayed some specific behaviours, we also used a manual selection of thresholds to group TFs (Figure 3A). We computed the mean AUC for the dense chromatin (QDA between 0 and 0.2) and open chromatin (QDA between 0.8 and 0.95) and the difference between these means. Using these values we classified TFs as: *(i)* accessibility independent factors (AIF) when both means are above 0.8 and the difference is lower than 0.1, *(ii)* accessibility dependent factors (ADF) when the difference is greater than 0.3 and the mean AUC in open chromatin is greater than 0.8, *(iii)* inaccessibility dependent factors (IDF) when the difference is greater than 0.3 and the mean AUC in dense chromatin is greater than 0.8, *(iv)* partial AIF/ADF when the difference is between 0.1 and 0.3 and the mean AUC in open chromatin is greater than 0.8, *(v)* partial AIF/IDF when the difference is between 0.1 and 0.3 and the mean AUC in dense chromatin is greater than 0.8 and *(vi)* poorly predicted if both means are below 0.65. TFs that did not meet any of these criteria were classified as other. The threshold of 0.8 AUC was selected based on the result from the K-means analysis by rounding the lowest AUC for the AIFs cluster, which was 0.79. In addition, we selected a threshold of 0.65 for poorly predicted TFs by rounding the average AUC for the *others* cluster in the K-means analysis, which was 0.64.

**Figure 2.**
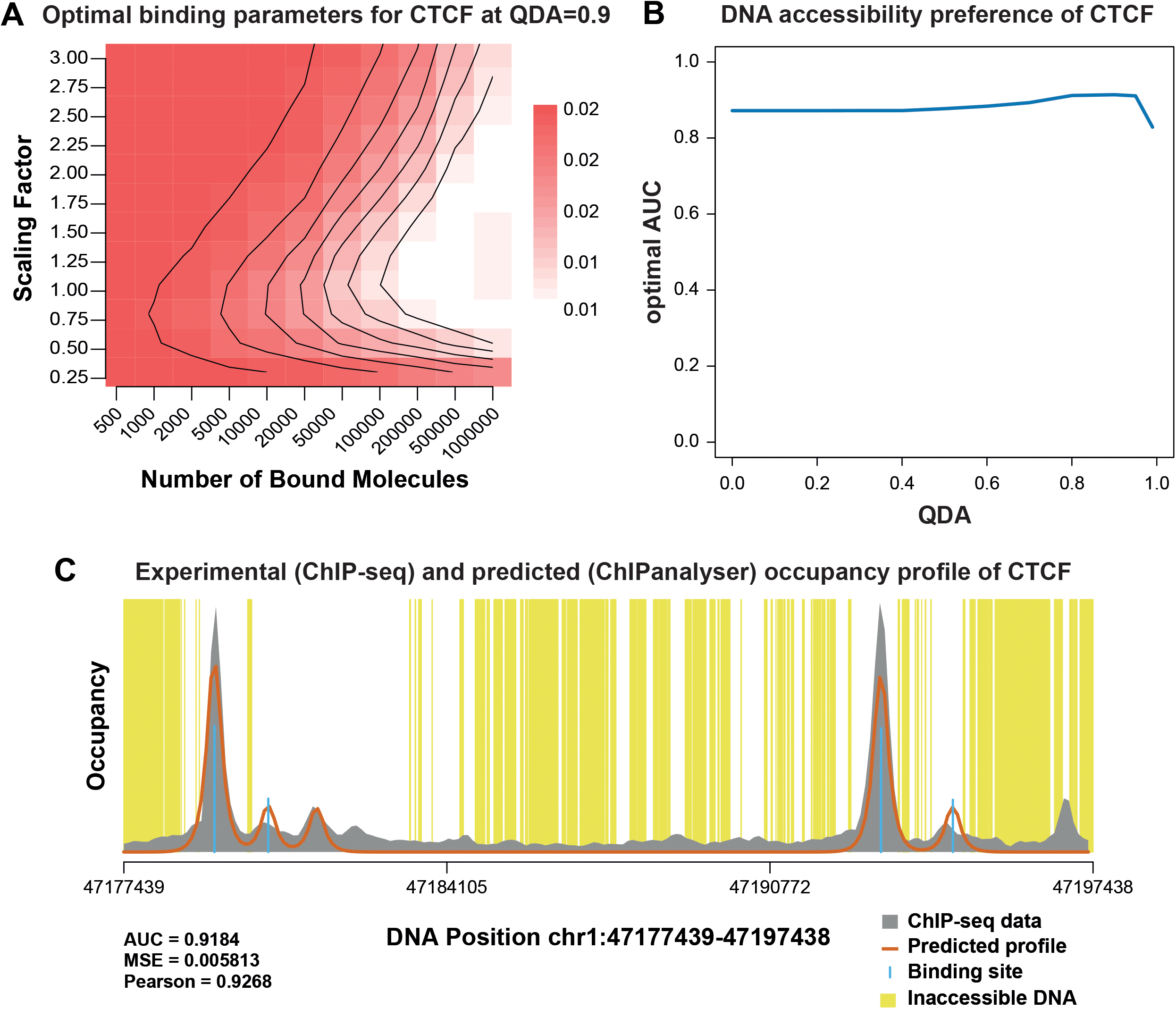
Model of CTCF binding. We plot the analysis of CTCF binding in K562 cells. (A) Heatmap showing the optimal range for the optimal QDA for CTCF (0.9). (B) AUC for the optimal parameters estimated for CTCF for all QDAs. (C) ChIP profile estimated with ChIPanalyser based on the optimal parameters. The grey shaded area represents the ChIP signal, the orange line represents the ChIPanalyser prediction of the ChIP signal, the blue lines represent occupancy at each locus, the yellow shaded areas represent closed chromatin, and the white shaded areas represent open chromatin.

**Figure 3.**
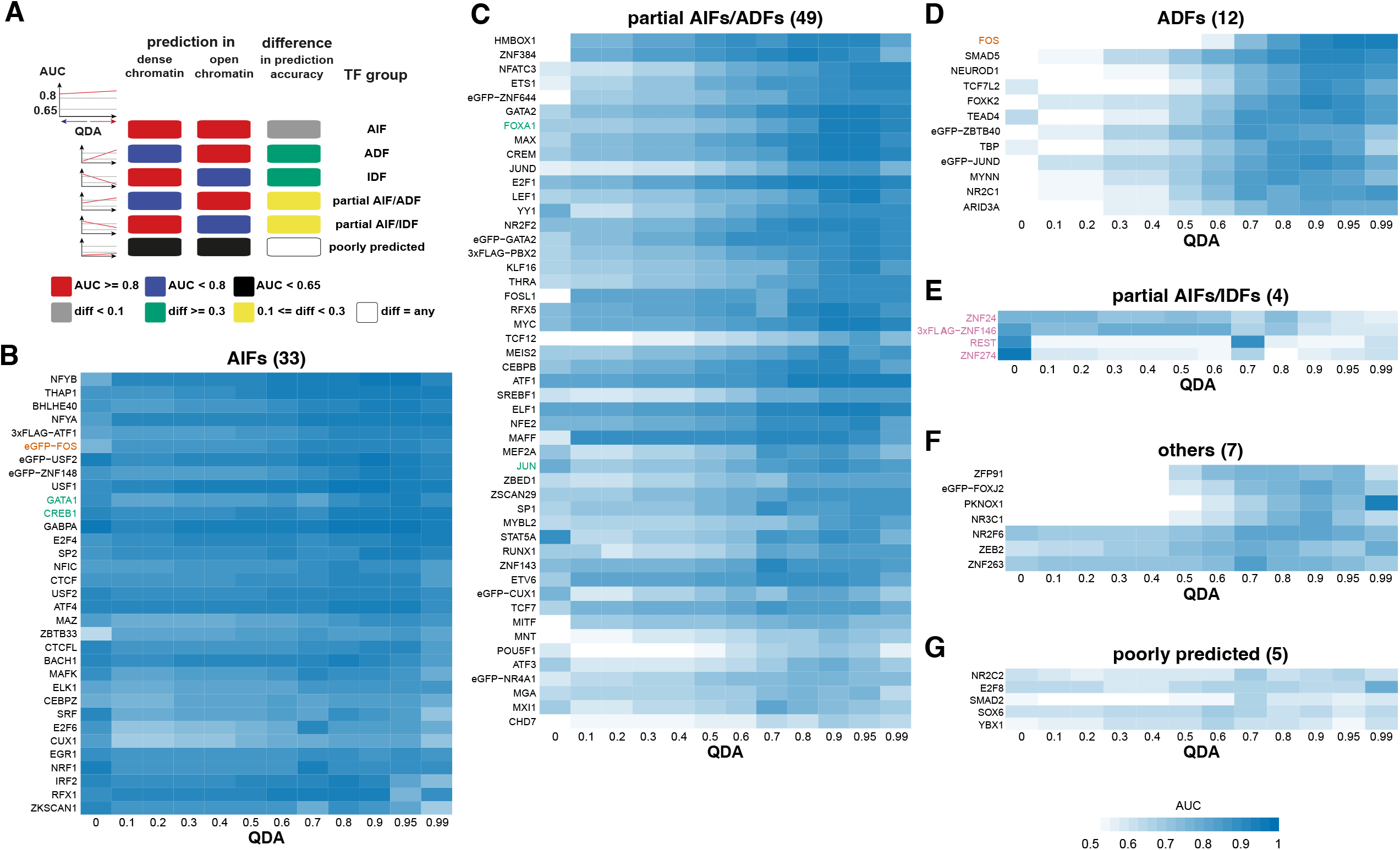
K562 classification of TFs. (A) We present the rules used to group the TFs in the different classes. (B-G) Heatmaps with the AUC for the optimal parameters estimated for each TF for all QDAs: (B) AIFs, (C) partial AIFs/ADFs, (D) ADFs, (E) partial AIFs/IDFs, (F) other and (G) poorly predicted. Each row represents a TF and each column an accessibility threshold (QDA value). The blue colour represents the AUC level for the corresponding QDA and TF. We mark with green and bold the TFs that were previously reported to act as pioneer TFs. For FOS, there are two different datasets leading to opposite results and that was highlighted by bold and orange colour of the text. Finally, we mark by purple and bold TFs that display a decrease in AUC with increasing the accessibility, which are potential Inaccessibility Dependent Factors (IDFs).

### Data access

All scripts used for pre-processing and further analysis can be accessed at https://github.com/nrzabet/human_TF_analysis.

## Results

### Modelling binding of human transcription factors

To investigate TF binding behaviour, we considered 110 TF ChIP-seq datasets as well as DNA accessibility data (DNaseI-seq) in K562 cells available from ENCODE (43, 44) (Figure S1 and Table S1). For each TF, we used a binding motif, in the form of position weight matrix (PWM), available from the JASPAR core database (56). To model the binding profiles of these TFs in K562 cells, we used ChIPanalyser (13, 37), a Bioconductor package that estimates ChIP-seq profiles based on four parameters: *(i)* binding motif of the TF in PWM format, *(ii)* DNA accessibility data, *(iii)* the number of TF molecules bound to the DNA (N) and *(iv)* a factor that modulates TF specificity (λ). The latter models the ability of a TF to discriminate between high and low affinity binding sites, where the affinity is estimated based on the PWM score. In particular, λ is inversely proportional to the capacity to differentiate between high and low affinity sites. In other words, a high λ means that there are more of the weaker binding sites, while low values for λ means that there are fewer but stronger binding sites for a TF. The number of molecules bound (N) and specificity factor (λ) are difficult to measure experimentally but can be estimated by fitting the model to ChIP-seq data and selecting the values that minimise the Mean Squared Error (MSE) between the predicted profile and the actual ChIP-seq profile (37) (Figure 1A). To run this analysis, the genome was tiled into 50 Kb bins and the model was then trained on 10 regions with strongest ChIP-seq signal. The top 10 regions of 50 Kb size contain the strongest peaks (true positives), but at the same time, they contain many regions that are not bound by the TF (true negatives). This provides an appropriate set of input data to train our model with sufficient true positives and true negatives (55). Following training, the estimated parameters were validated by computing the area under the curve (AUC) between the predicted profile and the ChIP-seq profile on the subsequent 50 regions with strongest ChIP-seq signal (37).

ChIPanalyser uses DNA accessibility data among other parameters to predict TF binding, and this provides the opportunity to investigate the role of DNA accessibility on the binding of TFs to the genome. We subset the DNA accessibility signal using a quantile vector into quantile density accessibility (QDAs) between 0-0.99 (37). In practice, this means that for each QDA, the model considers a percentage of the top ATAC-seq or DNaseI-seq signal regions as accessible, regardless of their actual accessibility scores (see Figure 1B-C and *Materials and Methods*).

To observe the preferences of the various TFs for chromatin accessibility, we ran the analysis for each QDA and used the AUC as a goodness of fit metric to estimate the model accuracy. If the AUC for the lower QDAs is high, it indicates that the TF can bind dense chromatin with high affinity. This is because the prediction accuracy remains high even though all or most of the genome is considered accessible to the TF by the model, including dense chromatin. In contrast, if the AUC is high only for the high values of QDAs, it indicates that the TF binds only in open chromatin, as only the most open regions are considered accessible to the TF by the model. We were able to identify three main classes of TFs based on their chromatin accessibility preference: Accessibility Independent Factors (AIF), Accessibility Dependent Factors (ADF) and Inaccessibility Dependent Factors (IDF) (Figure 1D). Each TF class has different DNA binding properties and impact on the surrounding chromatin, as illustrated in Figure 1D.

Figure 2 shows the analysis for CTCF in K562 cells. First, for each accessibility threshold value (QDA value), we fit the model to the ChIP-seq data over the training dataset and evaluate its performance over the validation dataset (Figure 2A-B). Figure 2B reveals that there are negligible differences between the performance of the model when assuming that CTCF can bind to all regions of the genome independent of their DNA accessibility levels (QDA=0) or when assuming that CTCF can bind only to the top 10% accessible regions of the genome (QDA=0.9). This means that CTCF binds to DNA in an accessibility independent way and, thus, can be classified as an AIF. Figure 2C shows an example comparison between the predicted CTCF profile and the ChIP-seq data, illustrating that the predictions are accurate. Similarly, Figure S2A shows two additional examples for other TFs (PBX2 and BACH1), while Figure S2B shows the distribution of AUC values across all TFs, which confirm that the predictions display high accuracy. A complete list of the estimated optimal parameters and AUC values for all TFs is available in Tables S2 and S3.

### Many human TFs are predicted to bind to the DNA at their strong binding regions independent of DNA accessibility

Following this workflow, we analysed the rest of the available ChIP-seq data in K562 cells. Figure S2 confirms that our model accurately fits the ChIP-seq data. More precisely, 95 out of 110 ChIP-seq datasets were modelled with high accuracy having an AUC of at least 0.8 and none have an AUC lower than 0.65 (Figure S2B). To group TFs with respect to their binding preference in open and dense chromatin, we first tried k-means clustering. We used K-means clustering to identify groups of TFs based on how well ChIPanalyser fits the ChIP-seq data (using AUC metric) assuming that the TF can bind in regions with different levels of accessibility. However, we were only able to detect two clusters despite observing multiple distinct behaviours when inspecting the AUC trends. For example, K-means clustering was not able to identify any IDFs. This is most likely due to having too few TFs exhibiting IDF behaviour in our dataset (see *Materials and Methods* and Figure S3). To evaluate if our original K-means analysis was too stringent, we also performed the K-means analysis with five clusters (Figure S4). This analysis resulted in grouping all IDFs together with poorly predicted ChIP-seq datasets and splitting the ADFs in two subgroups. Thus, we opted for a threshold-based approach to classify the TFs, where the values for the threshold selection were informed by the k-means clustering (see *Materials and Methods*). This analysis identified several classes of TFs with respect to their binding in open and dense chromatin (Figure 3A). Most importantly, the classification based on manual thresholds leads to similar results to K-means clusters (Figure S4F).

Unexpectedly, we found that many TFs display no preference for open or dense chromatin (33 AIFs), while some display a slight preference for open chromatin (49 partial AIFs/ADFs) (Figure 3B-C). CREB1, FOXA1, FOS, GATA1 and JUN were previously identified as pioneer factors (see Table S4) and, thus, their classification as AIFs or partial AIFs (green TFs in Figure 3B-C) is supported by the previous work. Only a small number of TFs displayed strong preference for open chromatin (12 ADFs) or moderate preference for dense chromatin (4 partial AIFs/IDFs) (Figure 3D-E). Seven TFs only partially met the criteria for either ADFs or AIFs and were classified as “other” (Figure 3F). Finally, five TFs (NR2C2, YBX1, SOX6, SMAD2 and E2F8) were not accurately predicted independently of the parameters that were used (Figure 3G), likely due to the quality of the ChIP-seq data or the PWM motif (Table S5).

It is worthwhile noting that FOS was classified as both an Accessibility Independent and Dependent Factor by our analysis (orange TFs Figure 3B, D and Figure S3B-C). Interestingly, the classification as an AIF was based on ChIP-seq data that was generated with an eGFP tagged version of FOS, while the ADF classification was based on the untagged version of FOS. One possibility is that, in the eGFP tagged experiment, the levels of FOS are higher than endogenous levels in K562 cells and at higher concentrations FOS could also bind in dense chromatin regions of the genome. This raises the possibility that a TF can act as an accessibility dependent factor in a cell line where it is expressed at low or medium levels and as an accessibility independent factor in cells where it is expressed a high or very high levels. In addition, it is known that eGFP can dimerise (57) and this could lead to a stronger recruitment of the eGFP tagged version of FOS compared to untagged version of FOS. One possibility is that homodimers of eGFP tagged version of FOS could have the capacity to bind dense chromatin.

It is also possible that some of the TFs identified as AIFs in our analysis bind in the same regions of the genome and our analysis identifies co-factors of TFs binding in high-occupancy target (HOT) regions (58) To investigate this, we looked for overlapping peaks of all TFs classified as AIF in K562 cells. Between the peaks used for model validation (based on which we performed TF classification), we found only negligible overlaps (Figure S5A), therefore the classification is not likely to be driven by TFs binding HOT regions. Furthermore, we also investigated overlaps among the top 1000 and top 5000 peaks and we observed that the extent of overlap increases with the addition of weaker peaks (Figure S5B-C), indicating that most of these TFs do not co-localise in the strongest binding regions in the genome, but do co-localise in weaker regions. The three main groups of TFs that co-localise are USF1 with USF2, CREB1 with ATF1 and SP2 with NFYA. TFs belonging to the same families, such as USF1 and USF2 or CREB1 and ATF1 are expected to share some lower affinity binding sites. Indeed, both pairs of TFs have similar binding motifs to each other. Furthermore, SP2 has been predicted to interact with NFY family members such as NFYB and NFYC (59). Therefore, the colocalisation of these TFs is most likely a result of similar function or recognition of similar low-affinity motifs and not due to many TFs binding in HOT regions.

Next, we used an alternative method to evaluate which TFs display strong binding in regions of the genome with low accessibility. In particular, we plotted the average ChIP-seq signal at 1000 regions with strongest and 1000 regions with lowest DNaseI-seq signal (Figures S6 and S7). We found that 9 AIFs (ATF4, CTCF, E2F6, EGR1, IRF2, MAFK, NFIC, NFYB and ZKSCAN1), 10 partial AIFs/ADFs (CEBPB, ELF1, HMBOX1, JUN, JUND, MAFF, MEIS2, NFE2, YY1 and ZNF384), 3 partial AIFs/IDFs (ZNF146, REST and ZNF274) and only 2 ADFs (ZBTB40 and MYNN) display at least similar ChIP-seq signal in dense chromatin regions as in open chromatin regions. Interesting, ZNF146 and REST showed stronger binding in dense chromatin than in open chromatin further supporting their classification as IDFs. Altogether, these results show that a large number of TFs that our analysis predicted to display DNA accessibility independent binding also show strong binding in dense chromatin.

In our analysis, we have considered all ChIP-seq peaks independent if they are promoter proximal or distal. Thus, we investigated if the classification of TFs as AIFs in our analysis has been impacted by their preference for proximal or distal binding to TSS. If open chromatin next to promoters were to dominate the signal (leading to classification of AIFs), then we should see that for all AIFs the majority of the ChIP-seq peaks used for the analysis are next to a TSS. In contrast, if dense chromatin was to dominate the signal (leading to classification of AIFs), then we should see that all AIFs have only a few peaks next to a TSS. What we see is a mixture (Figure S8B), which indicates that whether we train the model on TSS peaks or distal peaks does not influence the classification of TFs as AIFs or non-AIFs. We repeated the analysis by using only proximal or only distal ChIP-seq peaks for nine TFs that were selected since they were classified as AIFs, showed strong binding in dense chromatin (Figures S6A and S8A) and displayed different levels of TSS proximal and distal binding (Figure S8B). Our results showed only negligible differences in the AUCs of the nine TFs when considering either TSS proximal or TSS distal only binding, and TFs were consistently classified as (partial) AIFs (Figure S8D). Note when TFs that were classified as partial AIFs or other (ZKSCAN1, NFIC and MAFK), the analysis was performed on regions with weaker ChIP-seq signal (Figure S8C) because these TFs had few TSS proximal ChIP-seq peaks.

### Mouse TFs display similar behaviour to human TFs

Next, we performed a similar analysis for ChIP-seq datasets in mouse cell lines from the ENCODE project (43, 44). We obtained 60 ChIP-seq datasets covering 30 TFs (with a PWM motif in the JASPAR core database) in 8 cell lines (Figure S9 and Table S6). We also considered and additional study of MYOD1 from (60). Most of the ChIP-seq datasets (90%) were modelled with high accuracy (AUC >0.8); see Figure 4A. Depending on availability, we used DNase or ATAC-seq datasets as measures of DNA accessibility (see Table S7). For each ChIP-seq dataset, we computed the AUC for all QDA values (T able S8) and then used the threshold-based approach to group these TFs into the different classes (Figure 4B-H). Interestingly, we observed that while some TFs were classified in the same group for all cell lines, there were some that were classified in 2 or even 3 groups (Figure 4B). (60). Most of the ChIP-seq datasets (90%) were modelled with high accuracy (AUC >0.8); see Figure 4A.

**Figure 4.**
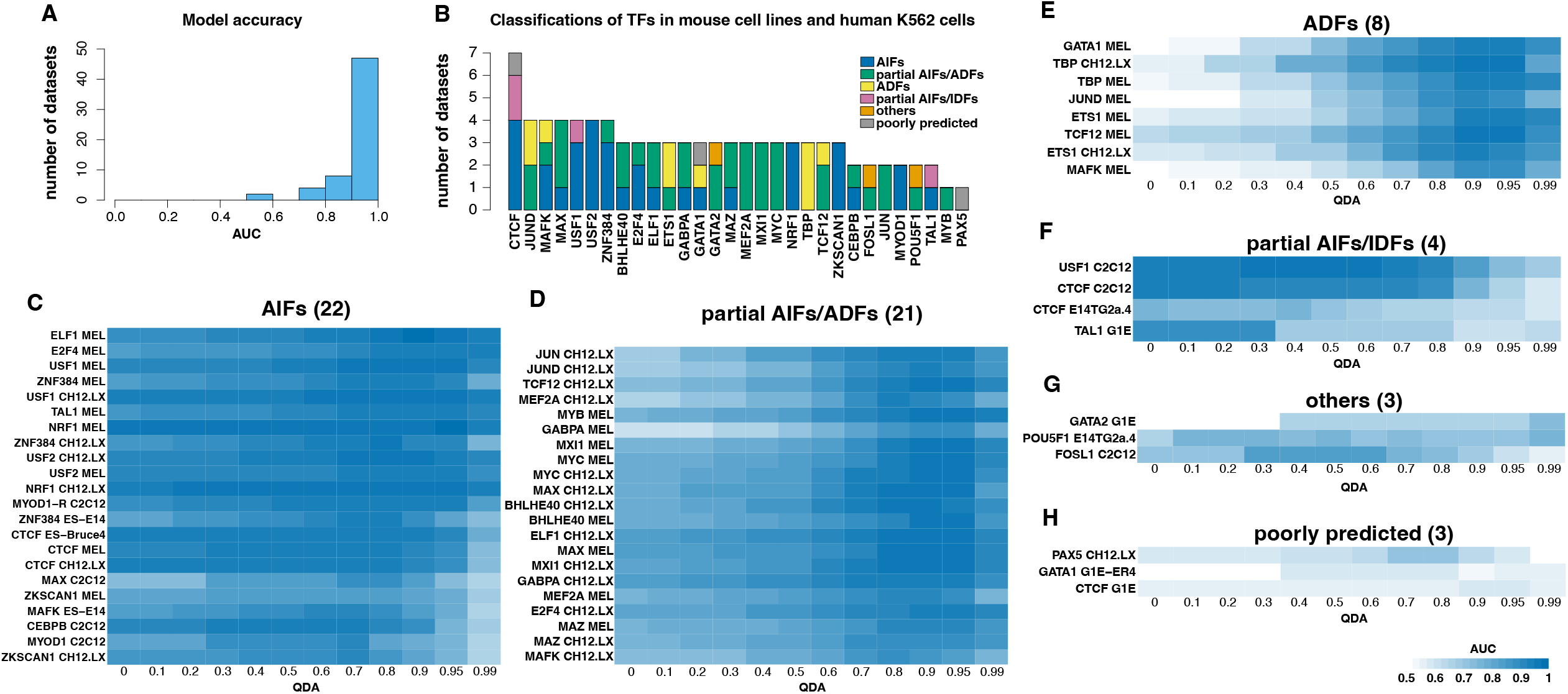
Classification of TFs in mouse cell lines. (A) Histogram with the AUC for the optimal parameters of the 60 TFs analysed in mouse cells. (B) Bar plot representing the different classifications for each TF in the mouse cell line. We also included the classification in human cells for the corresponding TF if that analysis was performed in K562 cells. (C-H) Heatmaps with the AUC for the optimal parameters estimated for each TF for all QDAs: (C) AIFs, (D) partial AIFs/ADFs, (E) ADFs, (F) partial AIFs/IDFs, (G) other and (H) poorly predicted. Each row represents a TF and each column an accessibility threshold (QDA value). The blue colour represents the AUC level for the corresponding QDA and TF.

Similarly, to our findings in the human cell line, CTCF was classified as an AIF in three of the mouse datasets. Interestingly, it was also classified as a partial IDF in two of the mouse datasets. This behaviour is in line with the literature surrounding CTCF, which suggests that its preference for chromatin state varies depending on context (34–36). Finally, CTCF was poorly predicted in one dataset, likely due to the quality of the data.

USF2 is the only TF that was identified in all four datasets (two in human and two in mouse) as an AIF suggesting that there is a high probability this TF binds in a DNA accessibility independent manner. USF1 displayed DNA accessibility independent binding characteristics in three datasets (two in mouse and one in human) and partial preference to dense chromatin in one dataset (mouse). These two TFs have been previously reported to bind heterochromatin (61), thus further supporting our findings. ZNF384 also displayed DNA accessibility independent binding characteristics in three datasets (mouse), but a partial preference to open chromatin in one dataset (human). NRF1 and ZKSCAN1 both showed consistent DNA accessibility independent binding in all three datasets. MYOD1, a previously reported pioneer TF (62–64) was identified in our study as displaying DNA accessibility independent binding in two different datasets. Only three TFs displayed consistent strong preference for open chromatin in at least two datasets, namely: TBP, ETS1 and JUND.

### Binding predictions are consistently better when using ATAC-seq and DNaseI-seq and are more accurate at regions with strong and medium ChIP signal

One surprising finding from our analysis in K562 cells is that many TFs are predicted to bind independently of DNA accessibility despite only a handful of TFs having been previously identified as pioneer TFs. This suggests that pioneer TFs are only a subset of AIFs and other types of TFs, such as co-factors of pioneers, bookmarking TFs or chromatin re-modelers also bind in an accessibility independent manner. However, our model is unable to distinguish between these types of factors. Moreover, the use of DNaseI-seq data might also be a potential source of bias and using different methods to estimate DNA accessibility could change some of these results. Similarly, we focussed our analysis at regions of DNA displaying the strongest ChIP signal and it is not clear if these results would remain the same when investigating regions of the genome with weaker signal.

To address these issues, we analysed the binding of eleven TFs in IMR90 cells and three TFs in HepG2 cells (Figure S10A-B and Table S9). First, we considered four different DNA accessibility measures (DNaseI-seq, ATAC-seq, MNase-seq and NOME-seq) in IMR90 and three (DNaseI-seq, ATAC-seq and MNase-seq) in HepG2; see Figure S10C-D and *Materials and Methods*. Secondly, we ran the validation analysis on 50 regions with strong, medium and weak ChIP-seq signal (see Table S9 and *Materials and Methods*). Our results showed that while most analyses resulted in a high prediction accuracy, there were significantly more cases compared to previous analyses where our model did not fit the data well (compare Figure 5A to Figure S2B and Figure 4A). To identify if there is a subgroup resulting in these lower accuracy models, we split the cases based on whether the validation ChIP signal was strong, medium or weak and found that most of the cases where our model did not fit the data well were at regions with weaker binding (Figure 5B). Next, we ran a similar analysis, but we split the data based on the DNA accessibility method. We found that while ATAC-seq and DNaseI-seq produced similar results, MNase-seq and NOMe-seq resulted in worse performance by our model (Figure 5C). Altogether, these results support that PWM, DNA sequence, binding specificity (λ), TF concentration and DNA accessibility can accurately explain observed binding profiles of TFs at regions of the genome displaying strong and medium binding strength, but only partially at regions displaying weaker binding. Furthermore, ATAC-seq and DNaseI-seq resulted in consistently better predictions than MNase-seq or NOMe-seq in our model, but this might be a reflection of the quality of those particular datasets (see Figure S10C-D and *Materials and Methods*).

**Figure 5.**
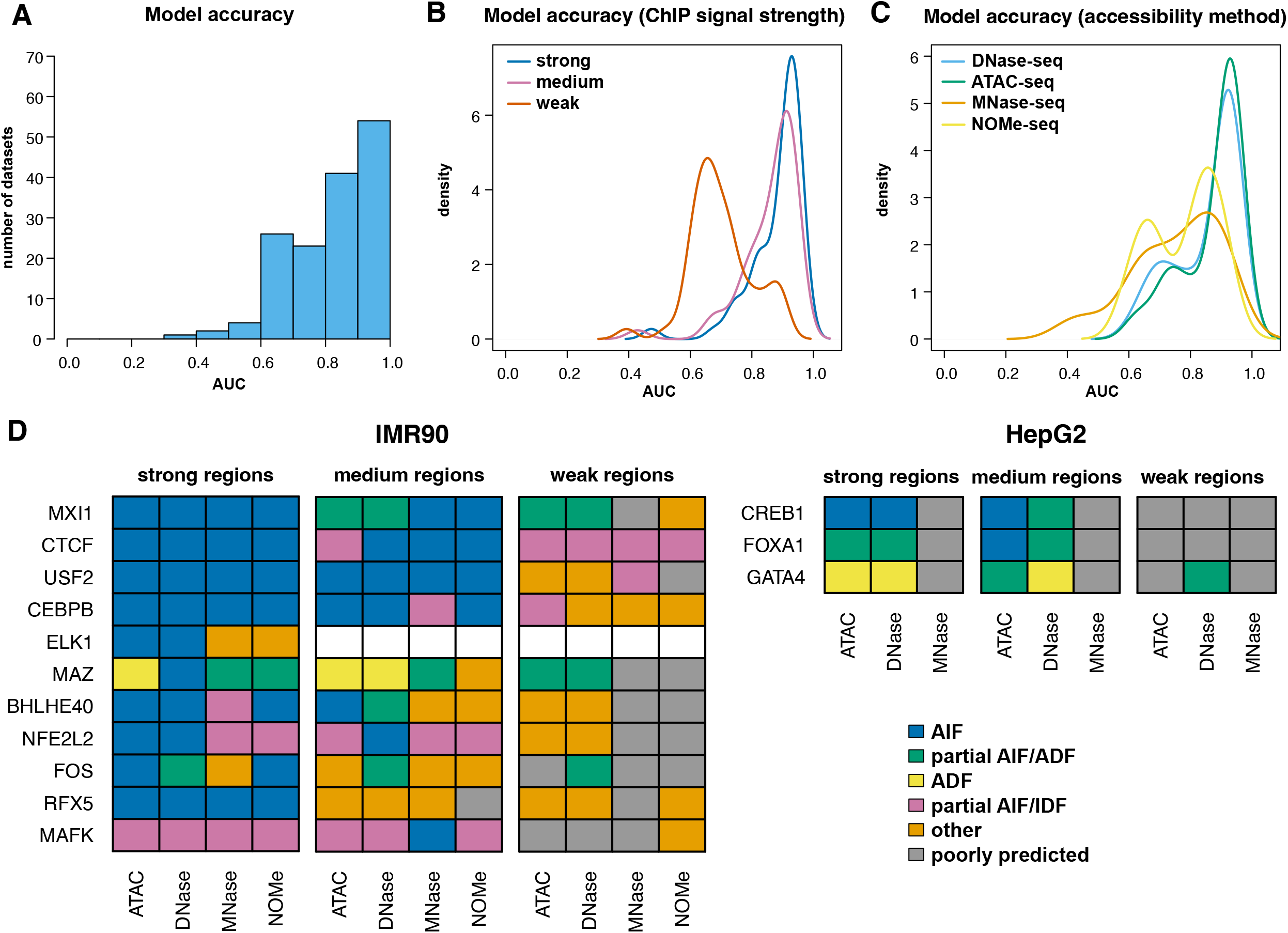
Effect of DNA accessibility method and ChIP-seq signal strength on TF classification. We consider 14 ChIP-seq datasets in IMR90 and HepG2 cells and investigated the effect of DNA accessibility method (DNaseI-seq, ATAC-seq, MNase-seq and NOME-seq in IMR90 and DNaseI-seq, ATAC-seq and MNase-seq in HepG2) and ChIP-seq signal strength (validation regions with strong, medium and weak ChIP-seq signals) on the classification of TFs. (A) Histogram with the AUC for the optimal parameters of all cases considered (13 TFs in two cell lines analysed with 3 or 4 DNA accessibility methods and strong, medium and weak binding genomic regions). (B) Density plot with the AUC for the optimal parameters of all combinations when the datasets are split based on ChIP-seq signal strength. We performed a Mann-Whitney U test and found that the differences in the means are statistically significant (p-value for strong compared to medium 0.046, strong compared to weak 2.46×10^-13^ and medium compared to weak 3.37×10^-11^). (C) Density plot with the AUC for the optimal parameters of all combinations when the datasets are split based on DNA accessibility method. We performed a Mann-Whitney U test and found that ATAC-seq and DNaseI-seq lead to similar results and these are different from MNase-seq and NOMe-seq (p-value for ATAC-seq compared to DNaseI-seq 0.32, MNase-seq compared to NOMe-seq 0.66, ATAC-seq compared to MNase-seq 3.03×10^-5^, ATAC-seq compared to NOMe-seq 3.52×10^-5^, DNaseI-seq compared to MNase-seq 2.97×10^-4^ and DNaseI-seq compared to NOMe-seq 5.15×10^-4^). (D) Classification of the 14 TFs in AIF, partial AIFs/ADFs, ADFs, partial AIFs/IDFs, other and poorly predicted.

In agreement with our findings in K562 cells and mouse cell lines, our analysis revealed that CTCF can be classified as both AIF and as partial IDF (Figure 5D, Figure S11 and Table S10). Interestingly, the latter is mainly associated with regions of weaker binding, while the former with strong and medium binding. Figure S12 shows an example where ChIPanalyser reproduces with high accuracy (AUC=0.95) the CTCF ChIP-seq data when assuming that it can bind anywhere independent of the level of DNA accessibility (QDA=0), but misses several peaks (AUC=0.5) when assuming that it can bind only to the top 5% accessible DNA (QDA=0.95). Similarly to CTCF, USF2, CEBPB and NFL2L2 act as AIFs at regions with stronger ChIP signal, but display IDF characteristics at weaker regions. RFX5 and MXI1 act as AIF, mainly at regions with strong binding.

FOXA1, CREB1 and FOS were previously identified as pioneer factors (Table S4) and were classified by our analysis as (partial) AIF (Figure 5D, Figure S11 and Table S10). Overall, we found that many TFs (nine out of fourteen) behave as AIFs, but preferentially at regions with stronger binding. At regions with weaker binding, they either behave as AIFs or our model cannot accurately capture their binding profile. MAFK is predicted most of the times to act as IDF and most likely to prefer dense chromatin. MAZ and GATA4 were identified to bind preferentially to open chromatin in regions with strong and medium binding.

One possible explanation for the observation that fewer TFs are classified as AIFs at regions with weaker binding is that they have weaker binding sites in those regions. To investigate this, we used PWMEnrich (65) to measure the strength of the binding sites located in peaks within strong, medium and weak binding regions. Our results showed that the majority of TFs (8 out 13) showed only small or negligible reduction in the strength of the binding sites located in medium or weaker binding regions (Figure S13). Nevertheless, one exception is CREB1, which displayed a large reduction in the strength of binding sites located in weaker binding regions compared to binding sites located in strong binding regions.

### Jun is predicted to bind before the chromatin is open in MCF10A cells upon Her2 overexpression

Recent work showed that HER2 overexpression in MCF10A cells resulted in a large number of regions gaining DNA accessibility and a small number losing DNA accessibility (66). One possibility is that these changes in DNA accessibility can be explained by changes in the levels of some AIF binding to those regions. HER2 overexpression also resulted in some TFs displaying changes in phosphorylation which could result in an increase or decrease of the number of bound TF molecules. We cross-referenced the TFs that were classified at least once as (partial) AIFs in our K562 experiment with the TFs that were shown to change phosphorylation upon HER2 overexpression and found seven TFs (ATF1, ETV6, JUN, JUND, MYC, NFATC3 and SRF). Using RNA-seq data, we estimated the number of bound molecules in MCF10A by multiplying the number of bound molecules in K562 cells with the ratio between the mRNA levels for the corresponding TF in the two cells, and kept λ the same as in K562 analysis (for each TF: N_MCF10A_=N_K562_ × mRNA _MCF10A_/mRNA _K562_ and λ _MCF10A_= λ _K562_). Rescaling the number of bound molecules by changes in RNA-seq between two cell types or two conditions was shown previously to generate results that reproduced ChIP-seq profiles with high accuracy (37). Then, for the HER2 overexpression experiment, we further rescaled the number of bound molecules based on the change in amounts of phosphorylated TFs (66).

Figure 6 shows the predicted binding levels at regions that gained and regions that lost DNA accessibility assuming that these seven TFs can bind independently of DNA accessibility levels. Interestingly, ETV6, JUN and SRF are predicted to bind strongest at these regions. SRF is the only TF that our model predicts to bind in an accessibility independent fashion and shows a noticeable increase in binding upon HER2 overexpression, but that happens at both regions that gained and regions that lost DNA accessibility (also see Figure S14).

**Figure 6.**
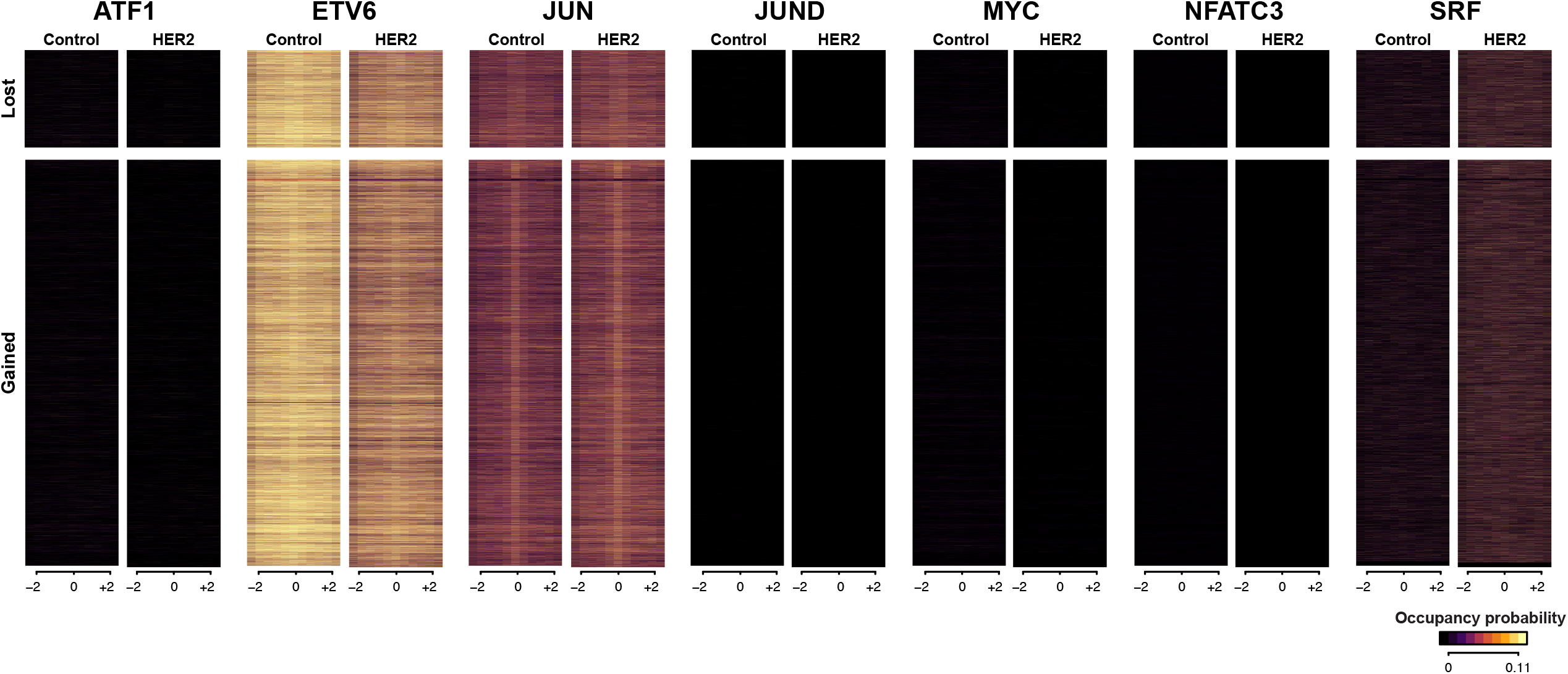
Prediction of TF binding in MCF10A cells upon HER2 overexpression. We considered the case of WT MCF10A cells and MCF10A cells were HER2 is over expressed. Overexpression of HER2 resulted in several regions gaining DNA accessibility and a few losing DNA accessibility. Heatmaps with the prediction of TF binding using ChIPanalyser for ATF1, ETV6, JUN, JUND, MYC, NFATC3 and SRF at regions that lost or gained DNA accessibility (±2 Kb) in control and HER2 over expression.

Our results show that there are negligible differences in the binding of JUN at regions that gained accessibility upon HER2 overexpression, suggesting that this TF is bound there before the regions become accessible. JUN is part of the AP-1 complex, which has been reported to have pioneer functions (see Table S4). Thus, while JUN can potentially bind inaccessible regions, there isn’t a subsequent opening of chromatin; instead, JUN appears to remain bound to closed chromatin. This is consistent with JUN being a bookmarking TF that binds to the DNA in dense chromatin and, when another co-factor is recruited, it could potentially open the chromatin.

## Discussion

Using our statistical thermodynamics framework, we systematically investigated the binding characteristics of a large number of human and mouse TFs in different cell lines and focussed our analysis on the relationship between TF binding and DNA accessibility. For several TFs, we also investigated the impact of using different DNA accessibility methods (ATAC-seq, DNaseI-seq, MNase and NOMe-seq) on our binding predictions in depth, to better understand potential sources of bias. In addition, we also compared the effects of modelling regions with stronger binding and regions that display weaker binding. Overall, we found that more TFs than previously reported do not display binding preference for open or dense chromatin (approximately one third), but this generally occurs at their strongest binding sites. Regions of the genome displaying weaker binding often are poorly modelled or show preference for either open or dense chromatin. In addition, we also provide evidence of DNA accessibility independent binding for TFs that have been previously reported to bind dense chromatin and explain their complex interaction with chromatin.

### Several transcription factors have no or limited preference for open or dense chromatin

A previous study proposed that approximately 16% of TFs display pioneer functions (42). Here, we find a higher proportion of TFs that bind chromatin independent of the DNA accessibility status (approximately one third). In addition to pioneers, AIFs also include bookmarking TFs (binding both open and dense chromatin) and co-factors of pioneers (being recruited by pioneer factors), which indicates that our estimates of the number of TFs that bind in an accessibility independent fashion are supported by this previous study.

Previous studies have reported that CTCF is a chromatin insulator (67). Nevertheless, CTCF also shows depletion of nucleosomes approximately 200 base pairs (bp) at the centre of its binding site (68). While the former suggests that CTCF prefers dense chromatin and does not open the chromatin, the latter indicates potential pioneer functions, where CTCF binds dense chromatin and opens it. In addition, it is reported that CTCF helps maintain the open chromatin state by steric hindrance of the DNA methylation machinery (69). Our results also showed that regions displaying weaker binding of CTCF could represent areas of the genome where CTCF acts as a potential insulator, while regions with strong binding reveal areas of the genome where CTCF might act as a pioneer factor. Altogether our results confirm that both functions are possible modes of action for CTCF.

Similarly to CTCF, CEBPB and NFE2L2 were identified as AIFs, mainly at stronger binding sites, and partial IDFs at weaker binding sites. CEBPB has been previously reported as having pioneer function and being able to maintain open chromatin (70), while NFE2L2 (NRF2) was reported as being a transcription activator (71). While our results support these previous findings, we also discovered a new role at weaker binding sites for these TFs. We propose that they preferentially bind the genome in denser chromatin and either maintain a closed chromatin state, or are unable to open the chromatin at those sites. This could be a consequence of the TF not being recruited in sufficient numbers to open the chromatin at weak sites, or it could indicate a bookmarking role, such as in the case of JUN.

USF1 and USF2 are members of the highly conserved basic-Helix-Loop-Helix-Leucine Zipper proteins (bHLH-LZ) that bind to the symmetrical DNA sequences called E-boxes (72). Previously, it has been hypothesised that USF1 and USF2 are able to bind heterochromatin and behave as pioneers (61). USF1 has been shown to form heterodimers that act as insulators, preventing the spread of heterochromatin (73). PLAG1 is one cofactor of USF2 that was suggested to display pioneer activity and enable USF2 binding, but the direct interaction was not proven (74, 75). Altogether, these previous findings support our results that USF2 binds as an AIF, and USF1 as an AIF or partial IDF.

Our analysis also identified NRF1, ZNF384 and BHLHE40 as AIF. NRF1 binding is methylation restricted in mouse embryonic stem cells, as it can be outcompeted by de novo DNA methylation. This suggests that methylation-sensitive TFs may rely on neighbouring pioneer TF binding to ensure a hypomethylated environment (76). ZNF384 was shown to interact with a variety of structural and regulatory proteins (vimentin, zyxin, PCBP1), but their individual roles in transcription regulation is not entirely clear (77). BHLHE40 was shown to induce binding-site directed DNA demethylation and hypothesised to have pioneer function (78, 79), which provides additional evidence for our classification as an AIF.

Many TFs in our analysis display accessibility independent binding, but this is mainly the case for regions of the genome for which they have the highest affinity. One question that arises is whether these TFs could truly bind independently of DNA accessibility. In our analysis, RFX5 was predicted to be an AIF only at regions with strong binding. Nevertheless, RFX5 was shown to displace nucleosomes, indicating that it has pioneer function (80). Altogether, these results support the fact that even if a TF is found to bind in a DNA accessibility independent way only at its strongest binding sites, it does not mean it cannot be a pioneer factor or that they are pioneer factors and not co-factors or bookmarking TFs.

FOXA1 is one of the best characterised pioneer TFs (28). In our analysis, FOXA1 was characterised as a partial AIF in K562 cells and in three scenarios in HepG2 (for regions with strong and medium binding) and only in one case as an AIF in HepG2 cells (at regions with medium binding when using ATAC-seq data) (Figures 3C and 5D). Interestingly, a previous study also reported that FOXA1 would have a reduced pioneer activity (42). Nevertheless, it was shown that despite its capacity to bind nucleosomes (18), its binding is chromatin context dependent (81). In other words, modifications of the histone tails might have an effect of FoxA1 capacity to bind nucleosomes and could explain why in some of our datasets and conditions we predict to display only partial accessibility independent binding.

JUN is part of the AP-1 complex, which has been proposed as a pioneer factor (82). Our analysis identified JUN as a partial AIF in K562 cells and mouse cell lines. Nevertheless, we predicted that JUN is already bound at regions that become accessible upon HER2 overexpression in MCF10A cells. This indicates that JUN is more likely a bookmarking TF that can bind in dense chromatin and requires co-factor(s) expressed upon HER2 overexpression to open the chromatin.

Finally, knockdown of MXI1 was shown to block chromatin condensation (83) suggesting that this TF could potentially act as an IDF. In our analysis MXI1 was consistently classified either as an AIF or as a partial AIF. While our analysis does not contradict the capacity of this TF to bind dense chromatin, it provides additional insights into its binding mechanism.

In our analysis, we did not consider the role of DNA methylation on TF binding. Previous work has shown that some TFs can distinguish between methylated and unmethylated DNA (84–88). Several previous *in vitro* studies (85, 89, 90) have observed that the binding of a large percentage of TFs seem to be affected by DNA methylation. However, in studies using cellular context, this is not the case and a recent paper showed that the function of 97% of enhancers is DNA methylation insensitive (87), indicating that binding of TFs *in vivo* is not DNA methylation dependent. This means that DNA methylation might impact the binding of only a small number of TFs considered here in native chromatin. Some of these TFs included in our analysis that preferentially bind unmethylated DNA include CTCF, GABPA, ELK1 and REST. The former three were classified as AIFs in our analysis and, for all three, ChIPanalyser predicts their ChIP-seq profiles accurately (AUC > 0.85) without considering DNA methylation. Figure S6 confirms that CTCF displays very strong binding in dense chromatin (with similar strength as in open chromatin) further supporting that its binding is not affected by DNA accessibility. In contrast, GABPA and CREB1 display significantly weaker binding at dense chromatin regions (Figure S6) indicating that they might not be capable of binding dense chromatin at least by themselves or that their binding is impacted by DNA methylation in dense chromatin. Overall, this shows that DNA methylation affects our results only marginally.

### Transcription factors that prefer dense or open chromatin

MAFK was found to be preferentially associated with dense chromatin in most cases and thus classified as an IDF. These results are supported by the fact that MAFK lacks a transactivating domain (91) and is mainly associated with heterochromatic parts of the genome (92).

We also identified four TFs with moderate preference for dense chromatin, namely: REST, ZNF274, ZNF24 and ZNF146. REST has been reported to be preferentially associated with silenced chromatin (93), ZNF274 is part of a complex that recruits H3K9me3 writers and, thus, is involved in heterochromatin establishment and maintenance (94), ZNF24 is associated with gene repression (95), while ZNF146 preferentially binds at silenced Line-1 elements (96). These previous reports support our classification of these TFs as IDFs.

TBP, ETS1, JUND, MAZ and GATA4 were TFs that were consistently identified to show binding preference for open chromatin. Interestingly, GATA4 has been reported to act as a pioneer factor (Table S4), but in our analysis, we found only partial AIF properties at regions with medium and weak binding (Figure 5D). This indicates that GATA4 function might be concentration- and chromatin context-dependent.

When investigating different datasets in the same cell type or in different cell types, we found that the classification of a TF may differ (e.g., Figure 4B). There are several possibilities that could explain this. One possibility is that the datasets have different qualities in terms of quality of the antibody or library and fragment size. Modelling datasets with different qualities can result in classification differences but selecting the most reproducible classification from biological replicates for the same TF can remove some of these biases. Alternatively, TFs will display different concentrations in different cell lines and, as seen in the analysis of JUN in K562, this can result in different preferences for dense chromatin. Finally, co-factors of TFs could be expressed in one cell type and not in another. This could help differentiate if a TF can open the chromatin by itself (it is modelled as an AIF in all cell types where it is expressed) or if it is also modelled as a bookmarking TF (modelled as an AIF in cell types where the pioneer co-factor is expressed and as an IDF where the co-factor is not expressed).

## Acknowledgements

The authors acknowledge the use of the High Performance Computing Facility at the University of Essex and would like to thank Stuart Newman for his support. The authors thank Zabet lab members for their useful discussions on the analysis.

