## Supplementary Material for "Identification of mammalian transcription factors that bind to inaccessible chromatin"

### Supplementary Table Captions

Table S1: *ENCODE ChIP-seq data in K562 cells.* Metadata for all ENCODE ChIP-seq datasets in K562 cells used in this study, including links to download the data and ENCODE identifiers.

Table S2: *Analysis results for ENCODE ChIP-seq data in K562 cells.* Information on pre-processing statistics (alignment rates and number of peaks), optimal parameters obtained from ChIPanalyser and classification based on the preference for DNA accessibility.

Table S3: *QDA data for ENCODE ChIP-seq data in K562 cells.* AUC for each QDA value for each TF analysed in K562 cells.

Table S4: *Summary of predicted/validated pioneer factors in humans in the literature.*

Table S5: *Poorly predicted TFs.* For each ChIP-seq that was poorly predicted in K562 cell line, we list the number of peaks called (low number of peaks can indicate poorer quality ChIP data), issues reported on ENCODE page for the corresponding dataset and information about the PWM motif used.

Table S6: *Pre-processing statistics for ENCODE ChIP-seq data in mouse cells.* For each TF and cell line analysed, the alignment rates and number of peaks are reported.

Table S7: *DNA accessibility data for mouse cell lines used in this study.*

Table S8: *QDA data for ENCODE ChIP-seq data in mouse cells.* AUC for each QDA value for each TF analysed in mouse cells.

Table S9: *Pre-processing statistics for ENCODE ChIP-seq data in IMR90 and HepG2 cells.* For each TF and cell line analysed, the alignment rates and number of peaks are reported together with the intervals for regions with strong, medium and weak binding.

Table S10: *QDA data for ENCODE ChIP-seq data in IMR90 and HepG2 cells.* AUC for each QDA value for each TF analysed in IMR90 and HepG2 cells.

### Supplementary Figures

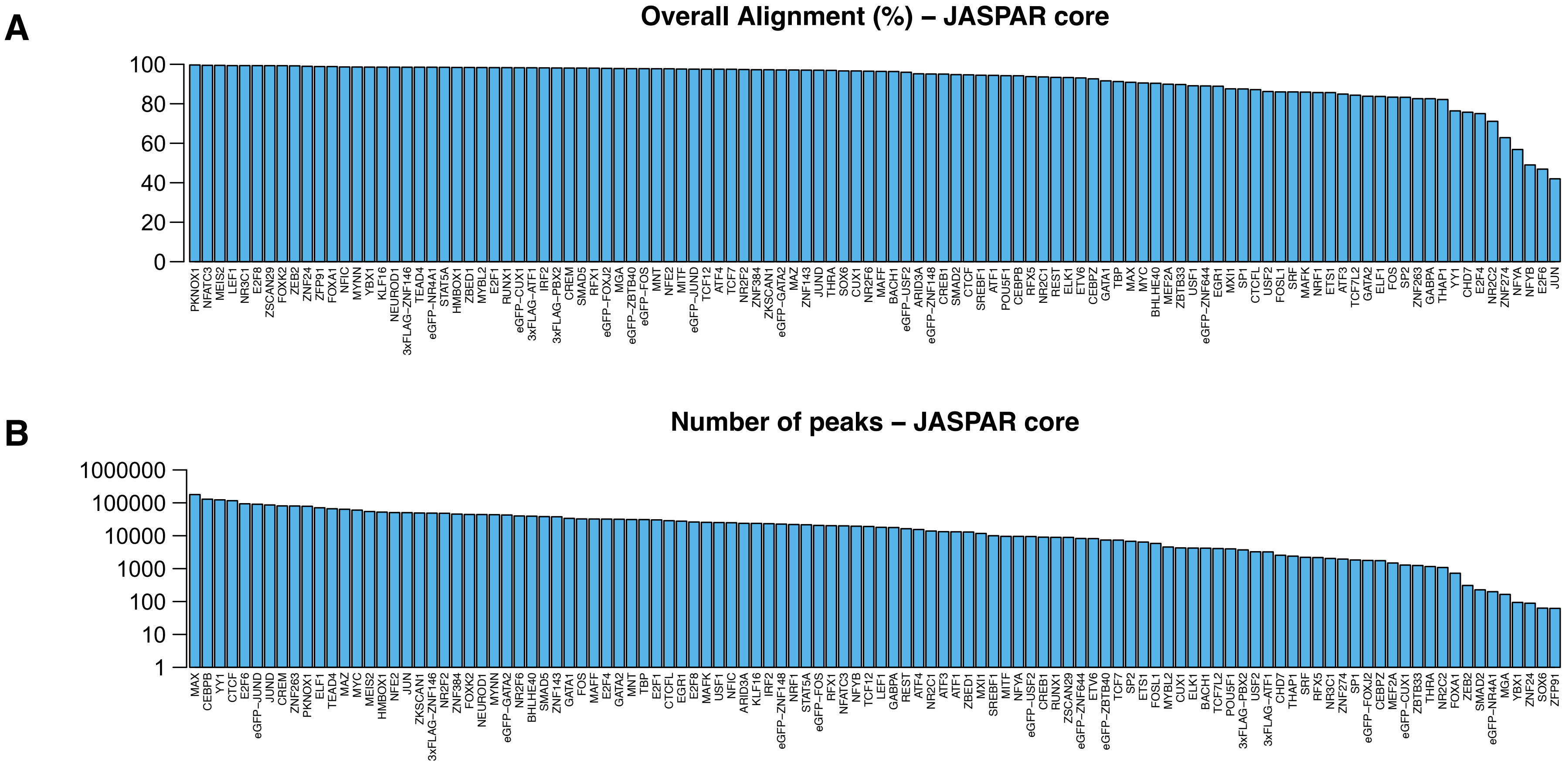


Figure S1: *K562 pre-processing statistics.* (*A*) The alignment rate of the ChIP-seq datasets for all analysed TFs in K562 cells. (*B*) The number of peaks detected for all analysed TFs in K562 cells.

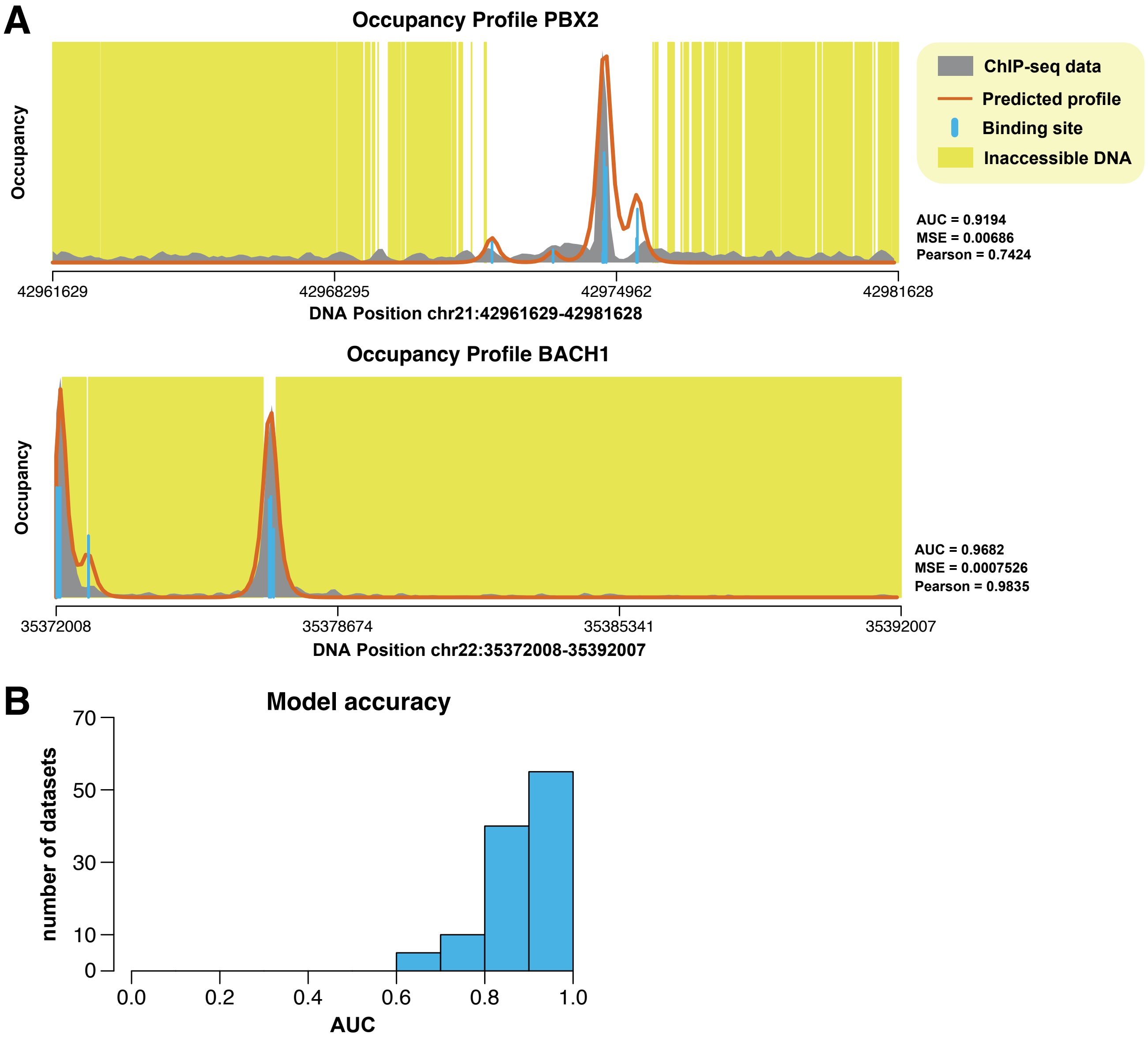


Figure S2: *K562 analysis statistics.* (*A*) ChIP profiles estimated with ChIPanalyser based on the optimal parameters for PBX2 and BACH1. (*B*) Histogram with the AUC for the optimal parameters of the 110 TFs analysed in K562 cells.

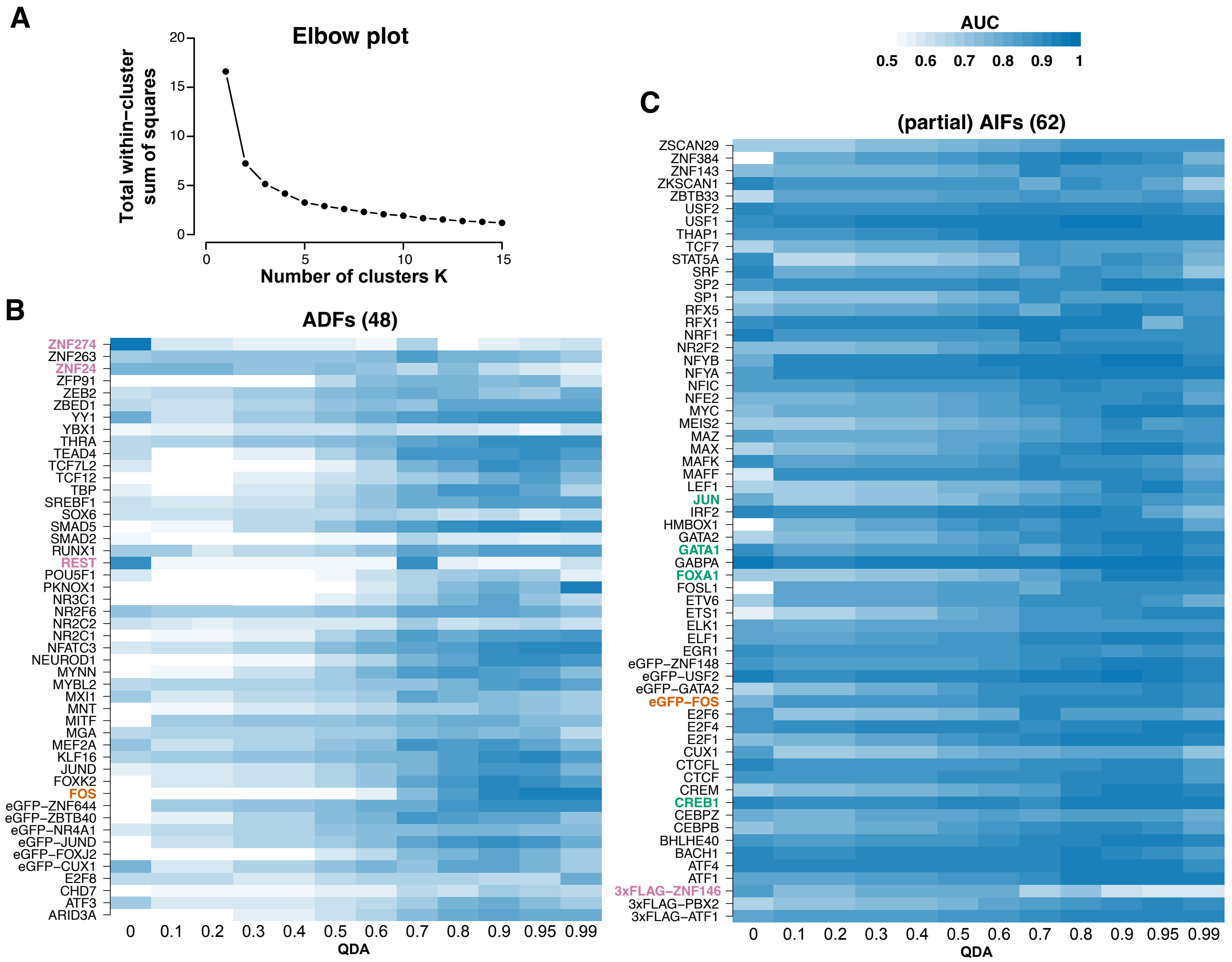


Figure S3: *K562 K-means clustering analysis.* (*A*) Elbow plot showing that the data can be split into two classes. ($B - C$) Heatmaps with the AUC for the optimal parameters estimated for each TF for all QDAs. Each row represents a TF and each column an accessibility threshold (QDA value). The blue colour represents the AUC level for the corresponding QDA and TF. We mark with green and bold the TFs that were previously reported to act as pioneer TFs. We mark by purple and bold TFs that display a decrease in AUC with increasing the accessibility, which are potential Inaccessibility Dependent Factors (IDFs).

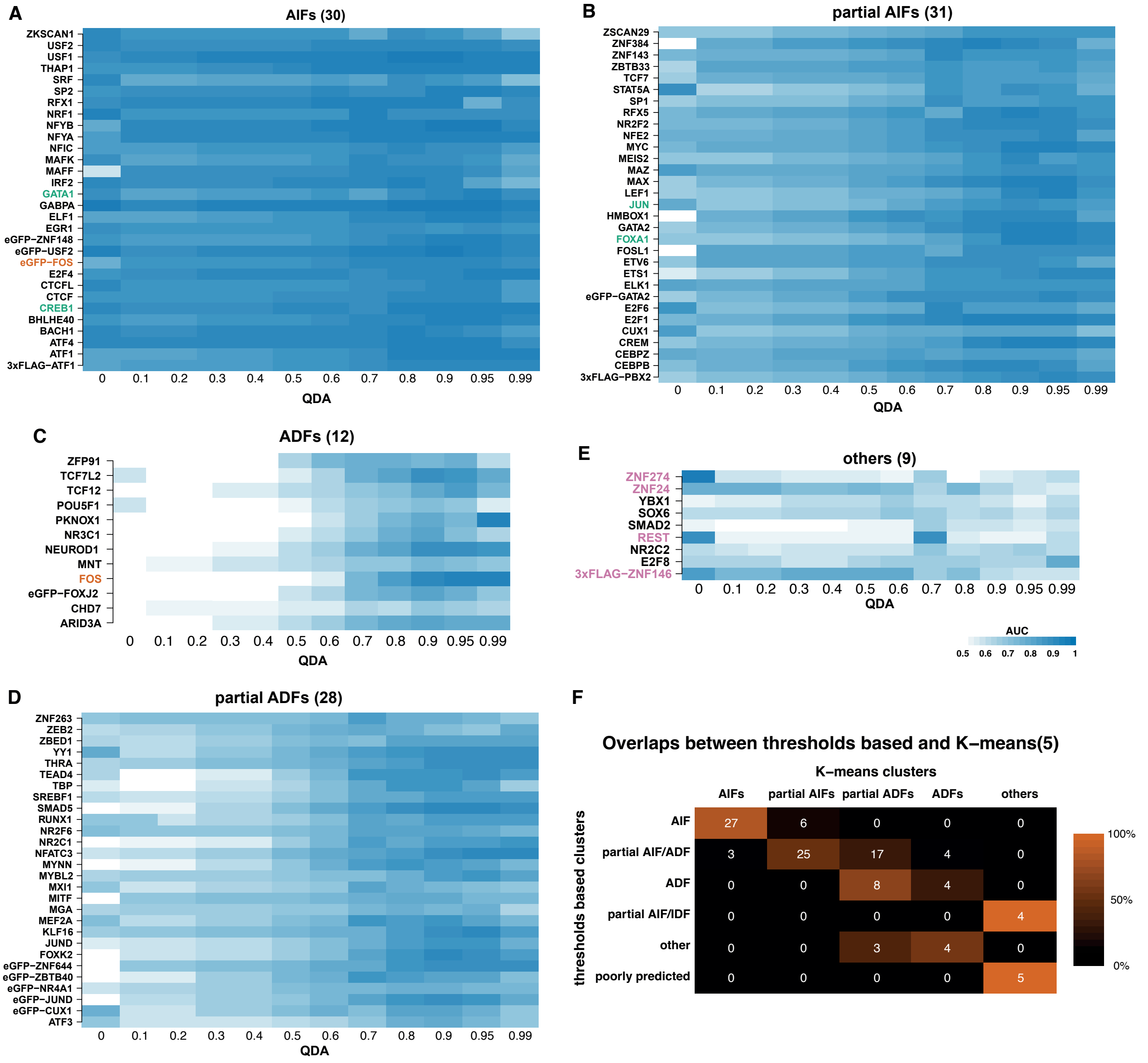


Figure S4: *K562 K-means clustering analysis with five clusters.* $(A-E)$ Heatmaps with the AUC for the optimal parameters estimated for each TF for all QDAs. Each row represents a TF and each column an accessibility threshold (QDA value). The blue colour represents the AUC level for the corresponding QDA and TF. We mark with green and bold the TFs that were previously reported to act as pioneer TFs. We mark by purple and bold TFs that display a decrease in AUC with increasing the accessibility, which are potential Inaccessibility Dependent Factors (IDFs). $(F)$ Comparison of the K-means and threshold based clustering. The analysis showed that the two approaches lead to similar results.

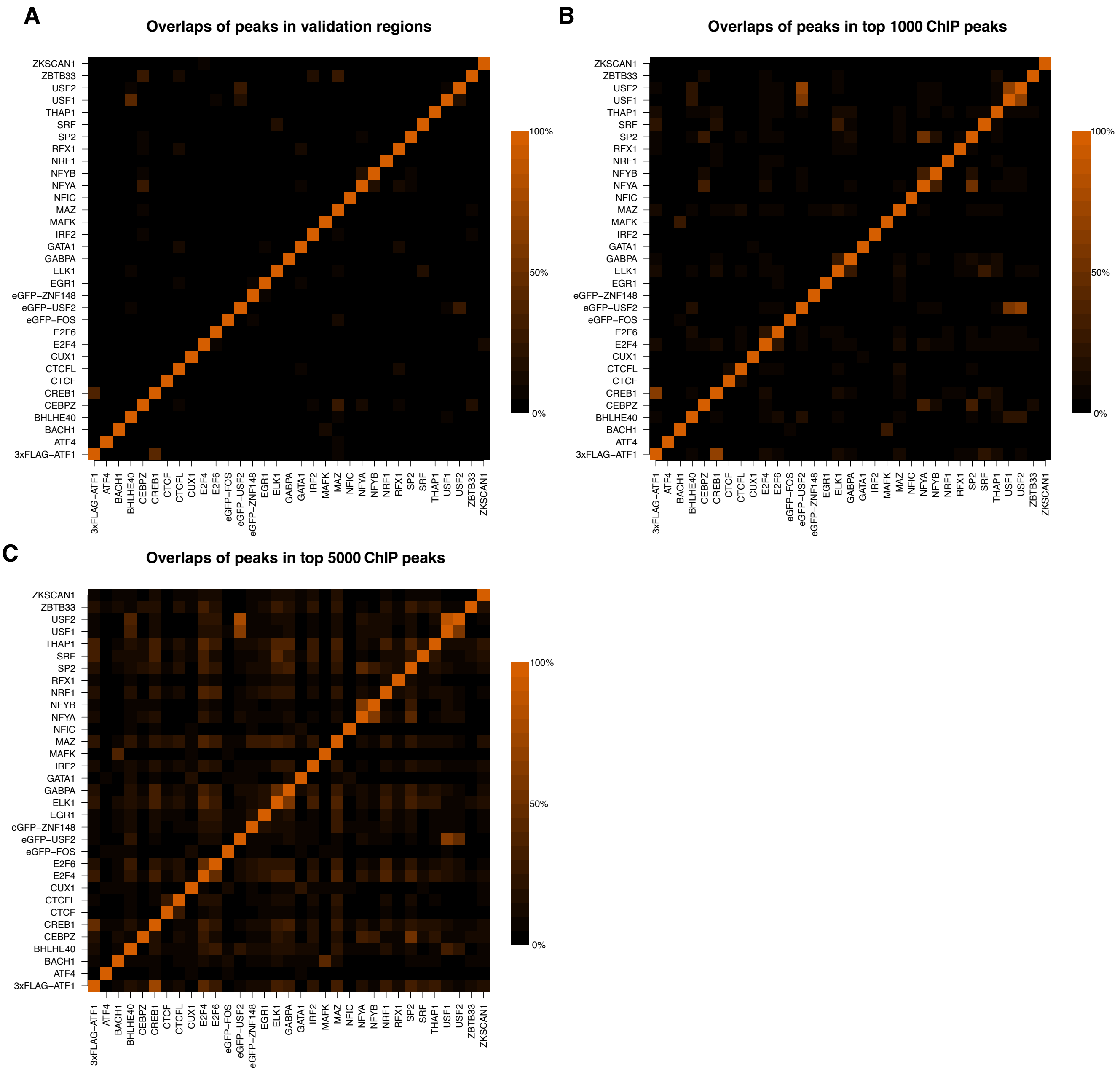


Figure S5: *Characterisation of ChIP peaks for AIFs in K562.* We computed the percentage of peaks for the ChIP-seq data on the rows that overlap with peaks ChIP-seq on the columns for: (*A*) ChIP-seq peaks in regions used for ChIPanalyser analysis, (*B*) top 1000 ChIP-seq peaks and (*C*) top 5000 ChIP-seq peaks.

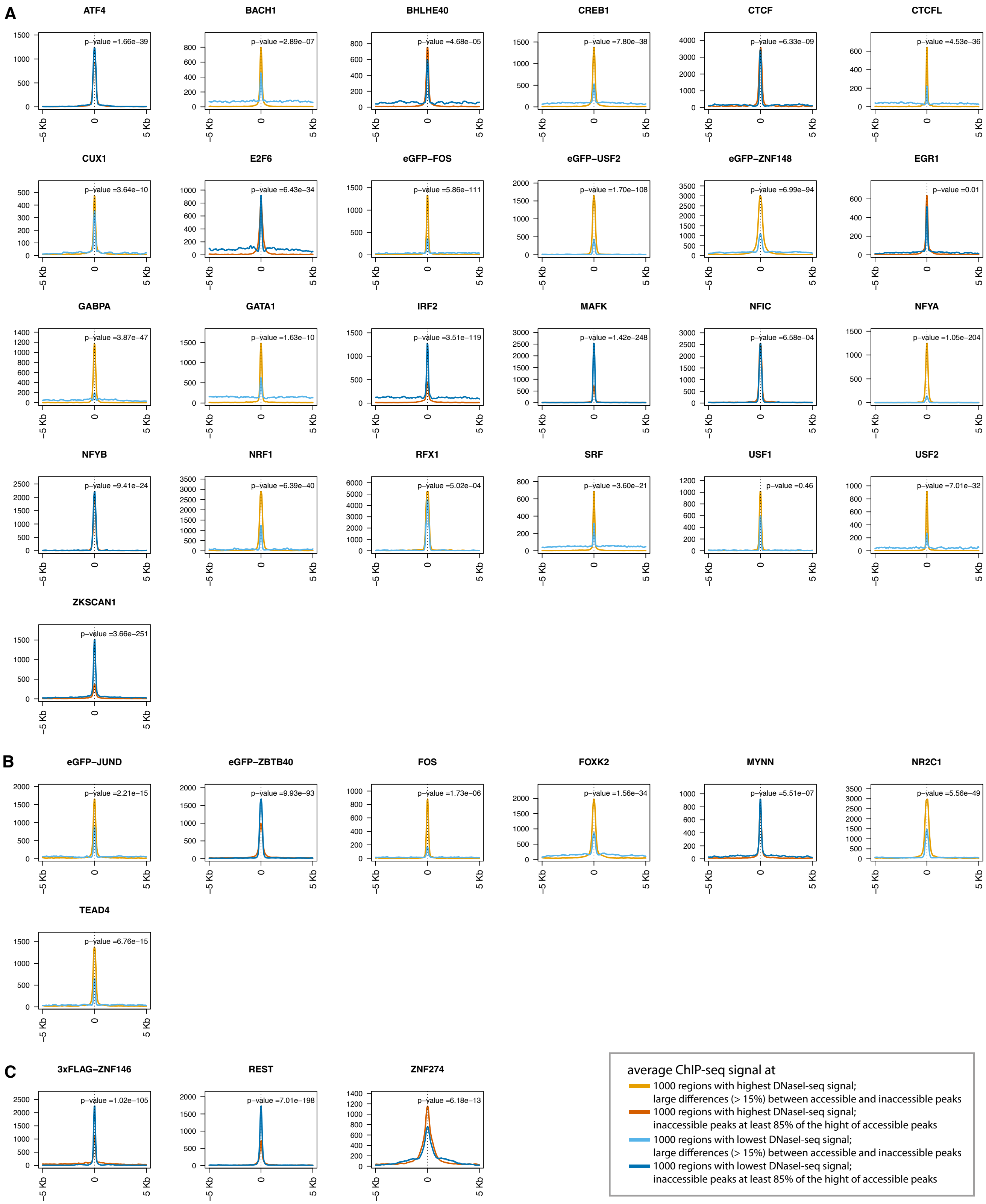


Figure S6: *Average ChIP-seq signal in K562 cells at most and least DNA accessible regions for AIFs, ADFs and partial AIFs/IDFs.* We considered the case of all TFs that have at least 2000 peaks and selected their ChIP-seq peaks in top (orange) and bottom (blue) 1000 accessible regions based on DNaseI-seq scores. We considered TFs that had a median ChIP-signal less than 10% of the maximum value in the 10 Kb region surrounding the centre of the peak. We used darker colours for TFs where the strength of the ChIP-seq signal in inaccessible regions is at least 85% of the ChIP-seq signal in accessible regions. and and lighter colours when the strength of the ChIP-seq signal in inaccessible regions is at most 85% of the ChIP-seq signal in accessible regions. We consider the case of (*A*) AIFs, (*B*) ADFs and (*C*) partial AIFs/IDFs

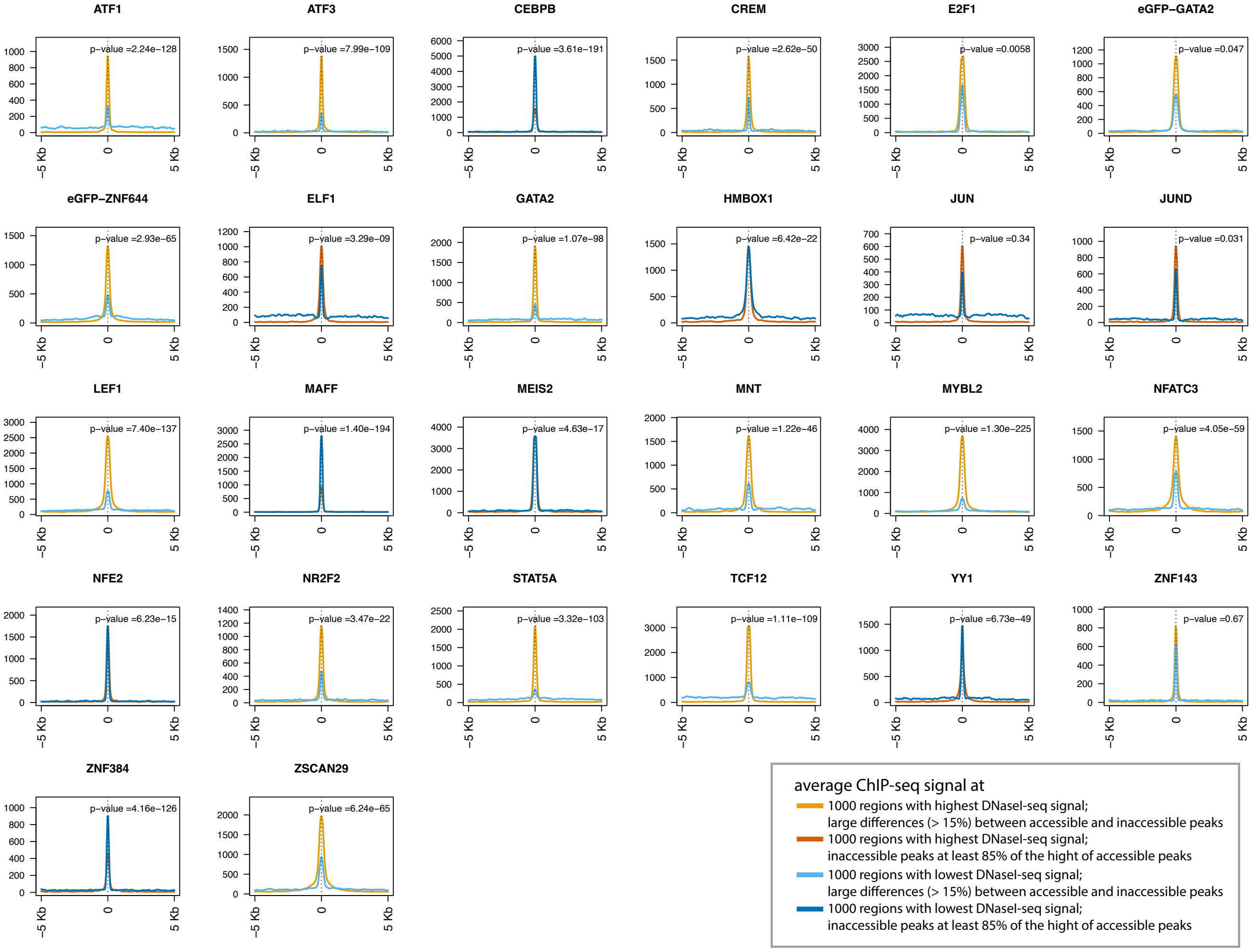


Figure S7: *Average ChIP-seq signal in K562 cells at most and least DNA accessible regions for partial AIFs/ADFs..* Same as in Figure S6 for partial AIFs/ADFs.

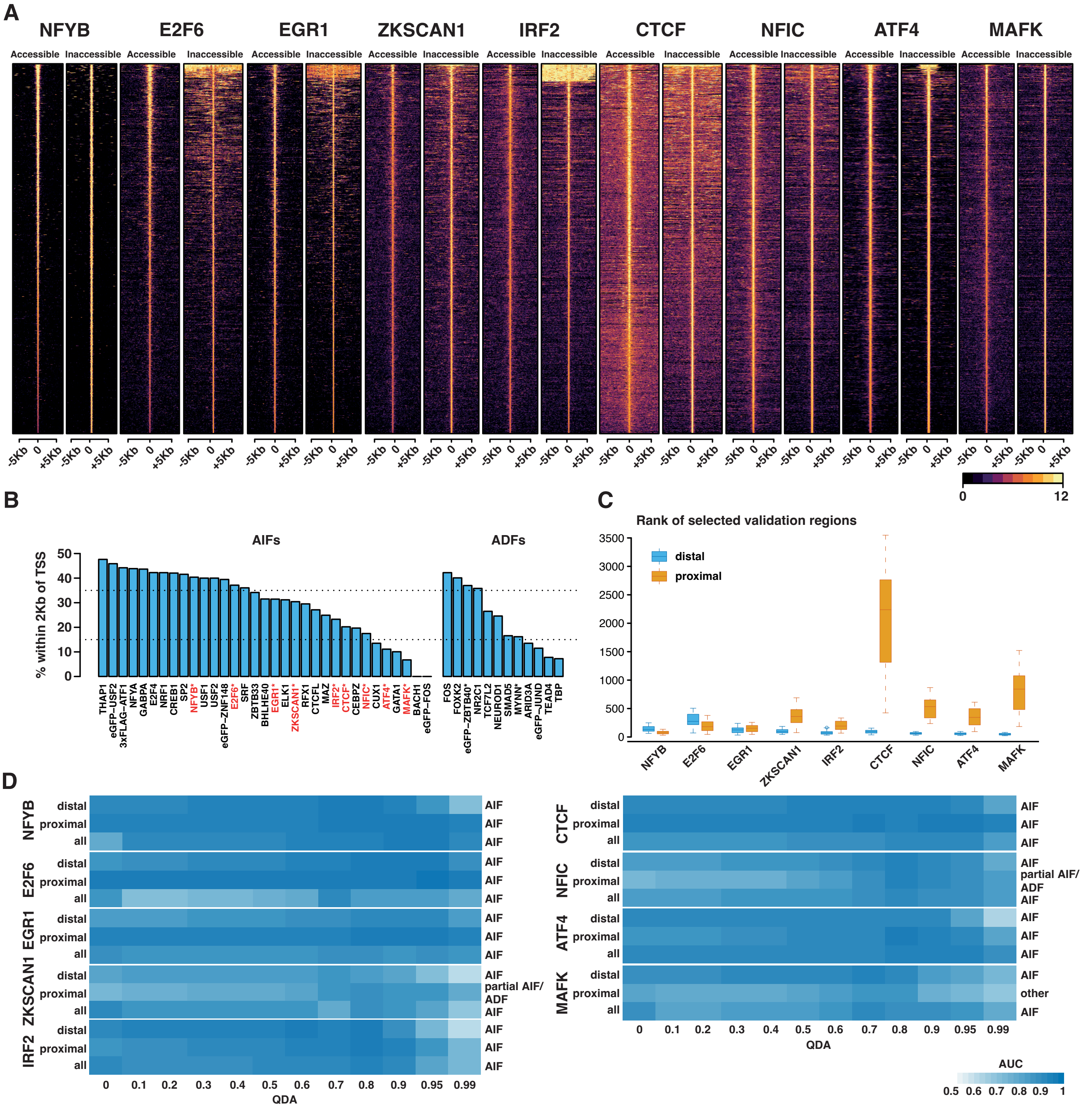


Figure S8: *Impact of proximity to TSS on the identification of AIFs.* (*A*) Heatmaps providing additional confirmation of strong binding in both accessible and inaccessible chromatin displayed by the nine AIFs identified in S6. Each line represents the $log_2$ transform of the ChIP-seq signal. (*B*) The percentage of peaks within 2 Kb of a TSS that were used in our analysis (top 60 regions with highest ChIP-seq signal) for each TF that was classified as AIFs and ADFs. We marked by * the TFs that showed strong binding in inaccessible DNA (see Figure S6). The dashed lines mark 35% and 15%. (*C*) The rank of regions ordered by ChIP-seq signal strength that were selected for validation when considering only TSS proximal (within 2 Kb of a TSS) or TSS distal (further than 2 Kb from a TSS) ChIP peaks. (*D*) Heatmaps with the AUC for optimal parameters estimated for each TF for all QDAs. We consider either all ChIP-seq peaks, proximal ones (within 2 Kb of a TSS) or distal ones (further than 2 Kb from a TSS) and each column an accessibility threshold (QDA value). The blue colour represents the AUC level for the corresponding QDA and TF.

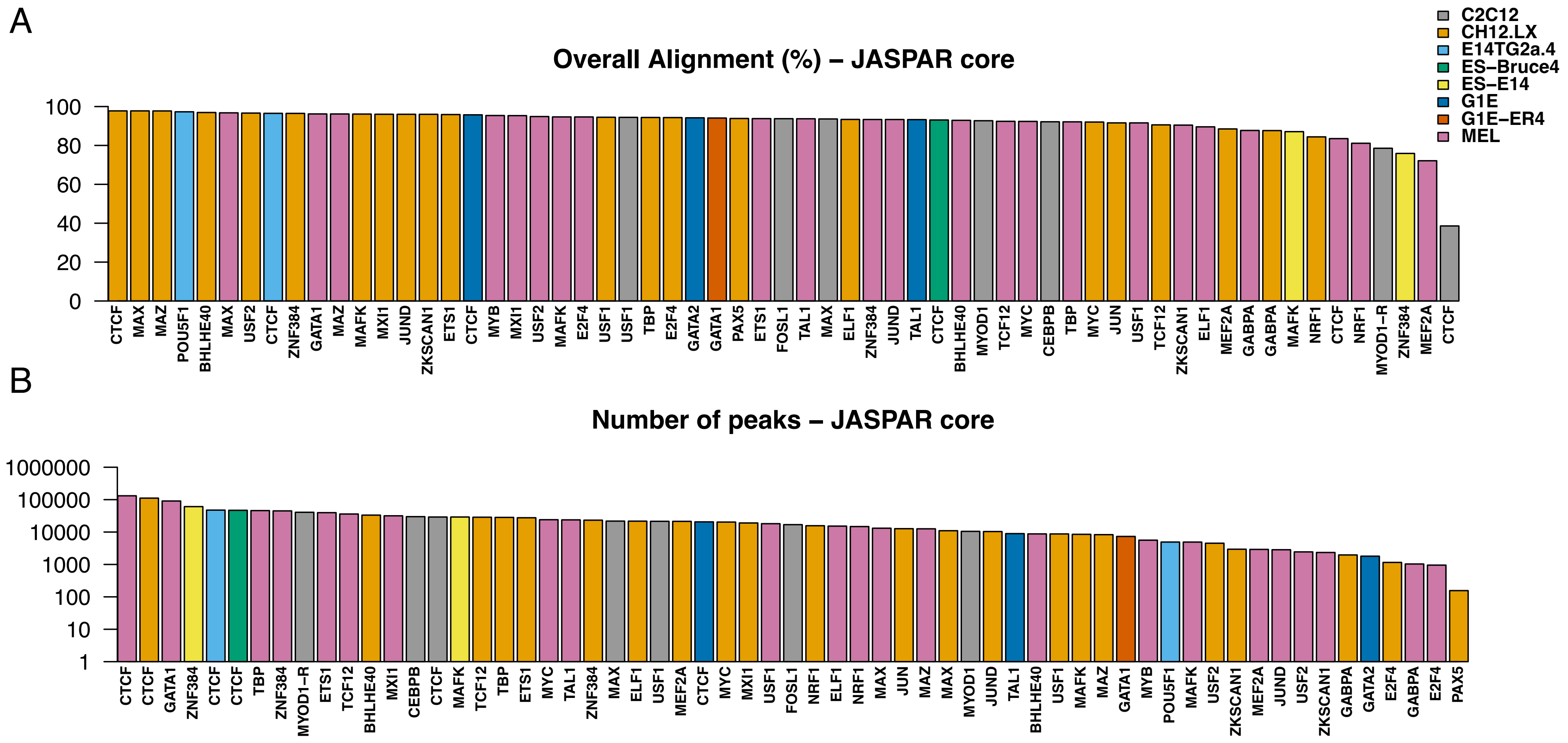


Figure S9: *Mouse ChIP-seq pre-processing statistics.* (*A*) The alignment rate of the ChIP-seq datasets for all analysed TFs in the mouse cell lines. (*B*) The number of peaks detected for all analysed TFs in the mouse cell lines. Each colour corresponds to one of the eight mouse cell lines (CH12.LX, E14TG2a.4, G1E, MEL, ES.E14, C2C12, G1E.ER4 and ES.Bruce4). MYOD1 represents the data from ENCODE, while MYOD1-R an additional dataset.

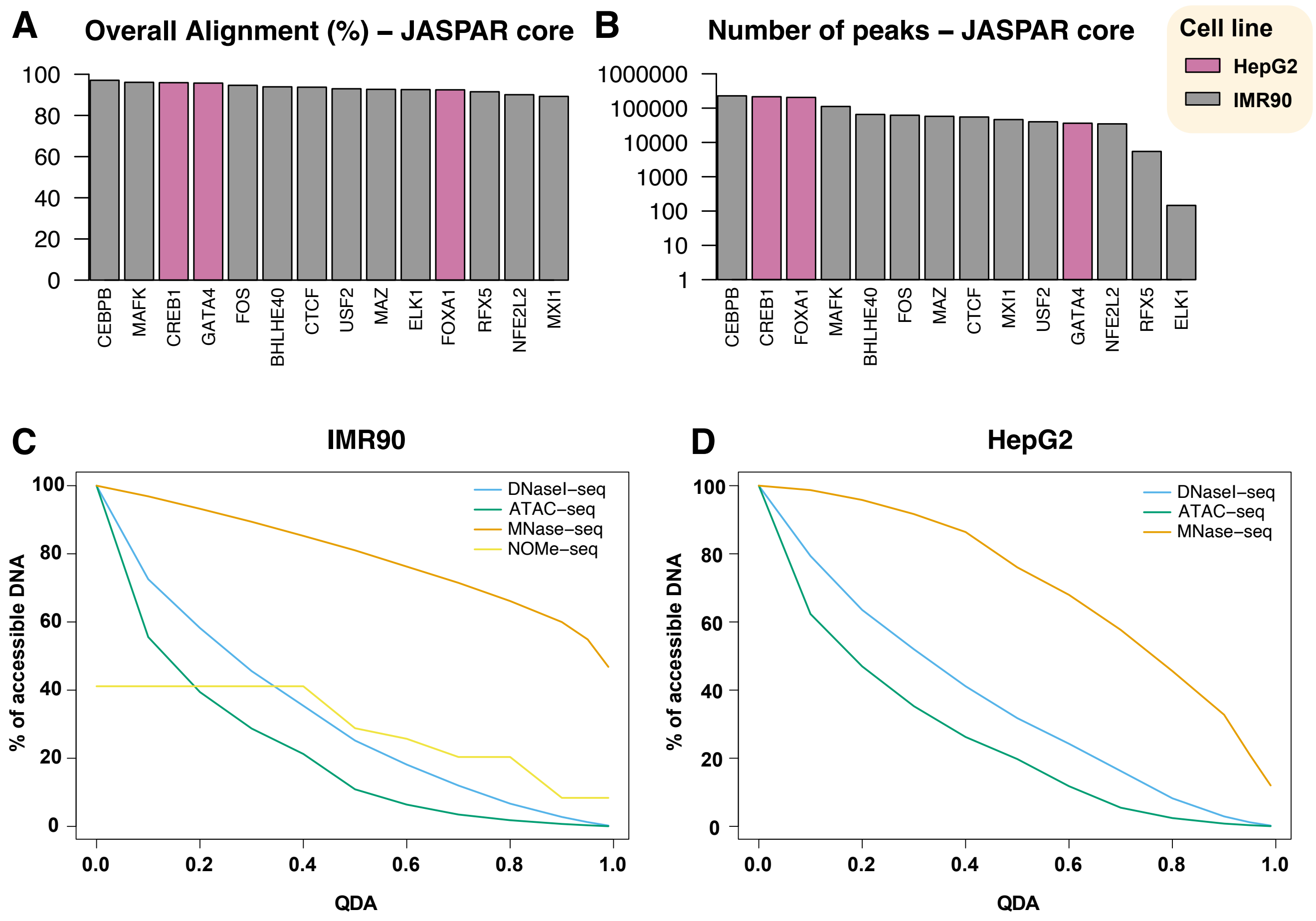


Figure S10: *IMR90 and HepG2 datasets pre-processing statistics.* (*A*) The alignment rate of the ChIP-seq datasets for all analysed TFs. (*B*) The number of peaks detected for all analysed TFs. (*C*) Percentage of accessible DNA for each QDA value in IMR90 cells. We considered four methods to measure DNA accessibility (DNaseI-seq, ATAC-seq, MNase-seq and NOME-seq). (*D*) Percentage of accessible DNA for each QDA value in HepG2 cells. We considered three methods to measure DNA accessibility (DNaseI-seq, ATAC-seq and MNase-seq).

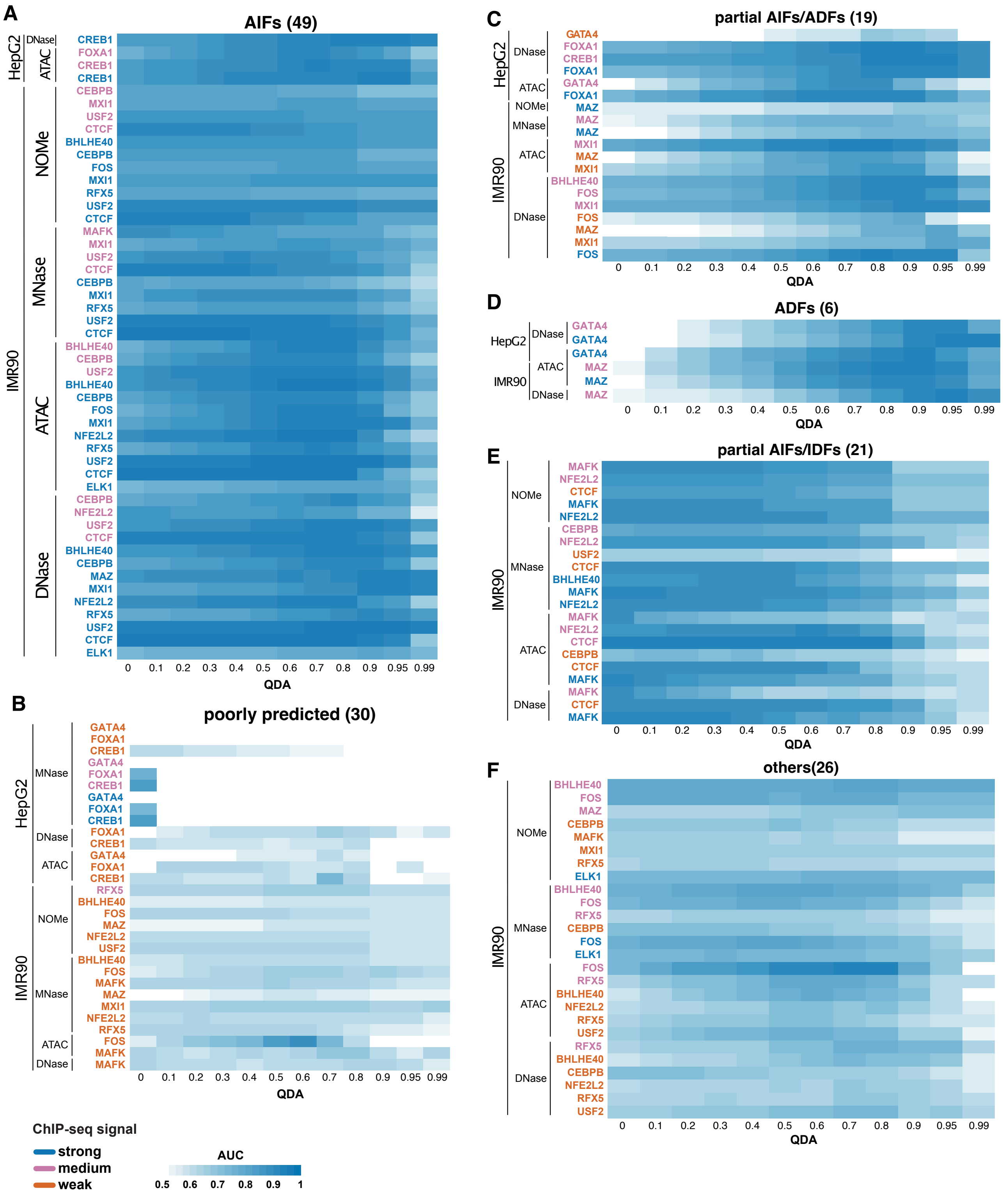


Figure S11: *Classification of TFs in IMR90 and HepG2 cell lines.* Heatmaps with the AUC for the optimal parameters estimated for each TF for all QDAs: (*A*) AIFs, (*B*) poorly predicted, (*C*) partial AIFs/ADFs, (*D*) ADFs, (*E*) partial AIFs/IDFs and (*F*) others. We considered the case of different DNA accessibility methods (DNaseI-seq, ATAC-seq, MNase-seq and NOME-seq) and running the validation step on 50 regions with strong, medium and weak ChIP-seq signals.

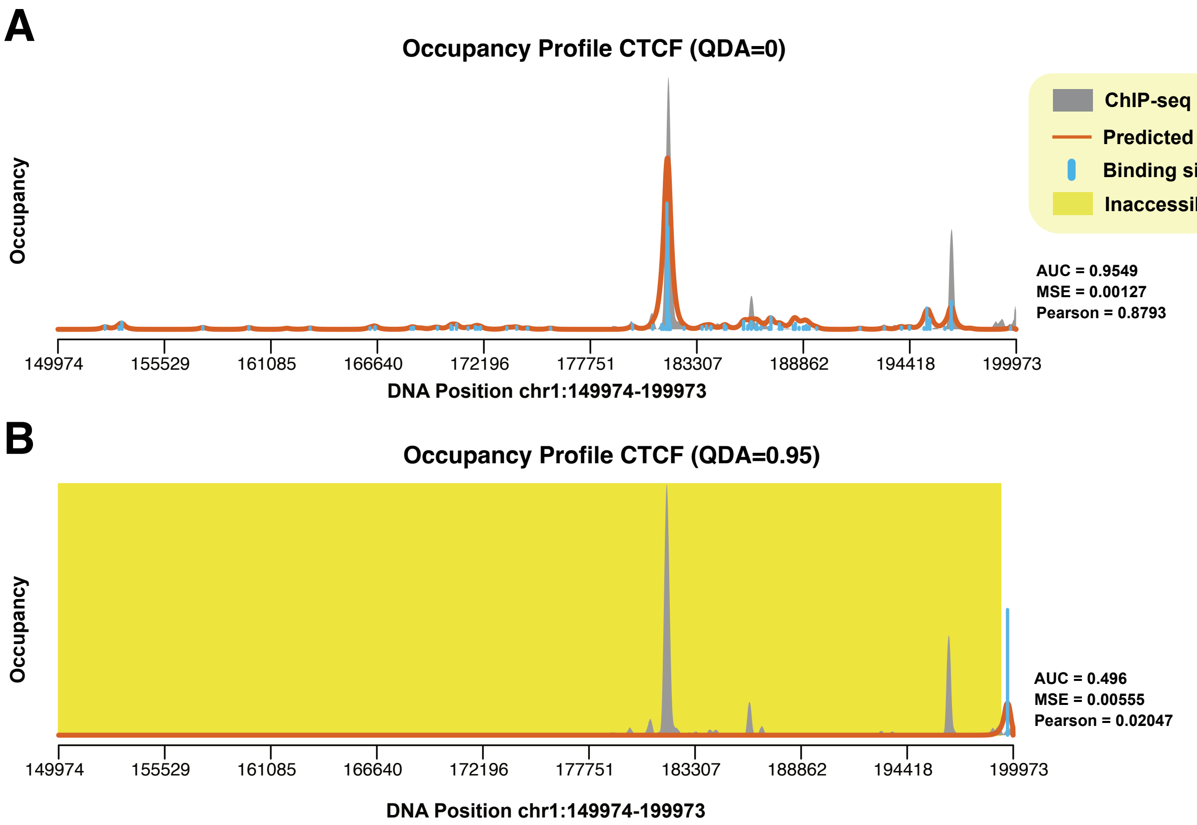


Figure S12: *CTCF binding in inaccessible DNA.* Example plots for CTCF binding in IMR90 cell line. (*A*) ChIP profiles estimated with ChIPanalyser based on the optimal parameters for CTCF assuming that it can bind anywhere, including in dense chromatin (QDA=0). (*B*) Same as in (*A*) , but assuming that CTCF can bind only top 5% accessible DNA (QDA=0.95).

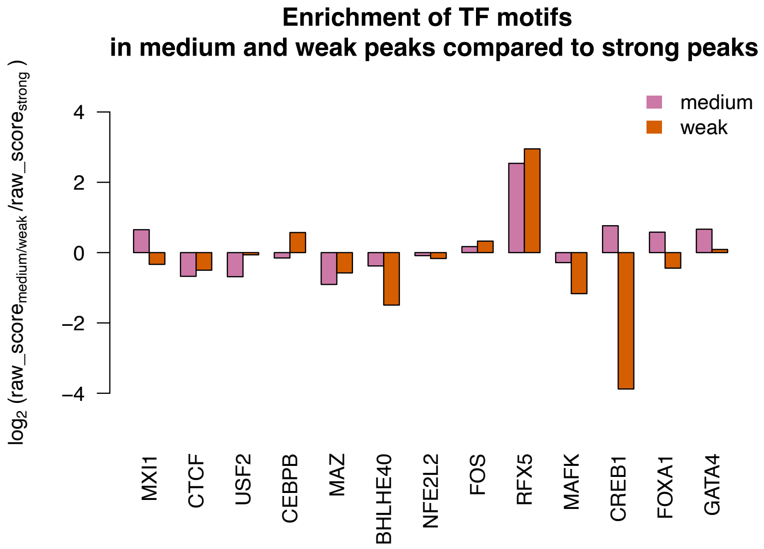


Figure S13: *Strength of binding for TFs at peaks in medium and weak regions relative to peaks in strong regions.* We computed the raw score for the TF binding estimated by PWMEnrich and then performed the $log_2$ ratio of the raw score at peaks in medium (magenta) or weak regions (orange) compared to the raw score at peaks in strong regions.

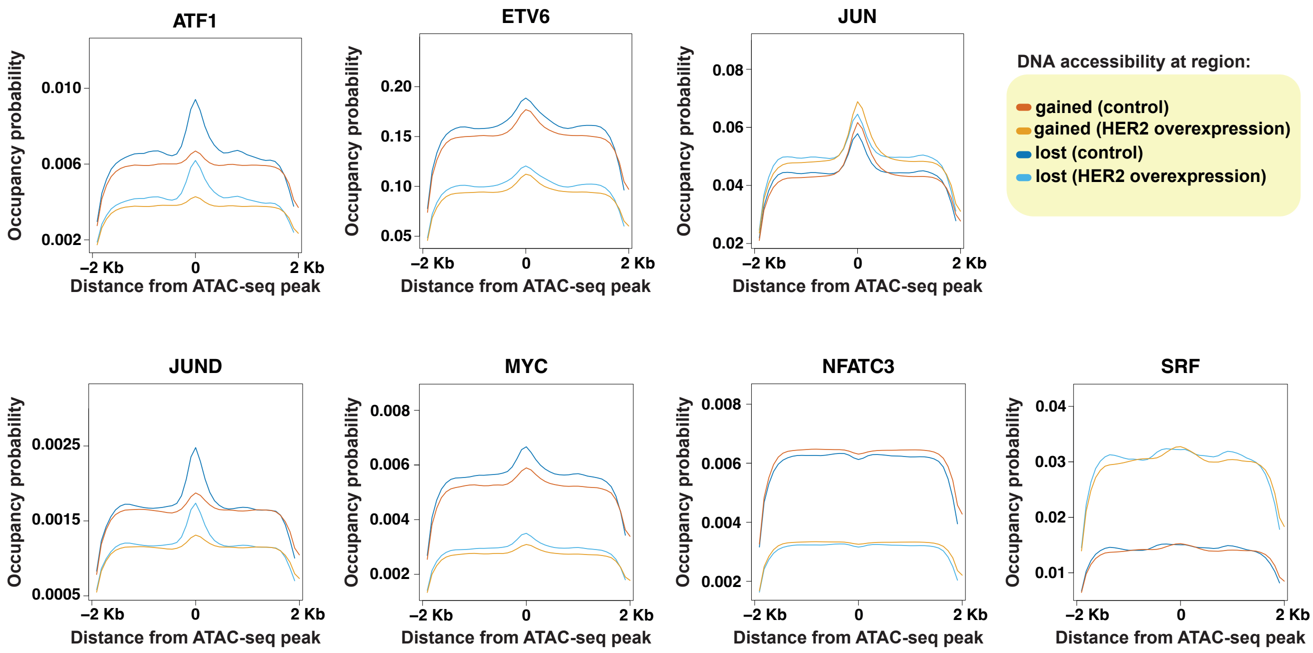


Figure S14: *Average profiles of TF binding in MCF10A cells upon HER2 overexpression.* Average profiles of the prediction of TF binding using ChIPanalyser for ATF1, ETV6, JUN, JUND, MYC, NFATC3 and SRF at regions that lost or gained DNA accessibility in control and HER2 over expression.
