## Supplementary material for "Identification of mammalian transcription factors that bind to inaccessible chromatin": Table S4

**Table S4.** Summary of predicted/validated pioneer factors in humans in the literature.

| TF name | Reference |
| --- | --- |
| FoxA1/2 | reviewed in (1) |
| Oct3/4, Pou5f3 | reviewed in (1) |
| Sox2 | reviewed in (1) |
| Klf4 | reviewed in (1) |
| Ascl1 | reviewed in (1) |
| Pax7 | reviewed in (1) |
| PU.1 | reviewed in (1) |
| GATA4 | reviewed in (1) |
| GATA1 | reviewed in (1) |
| CLOCK:BMAL1 | reviewed in (1) |
| P53 | reviewed in (1) |
| Pbx1 | reviewed in (2) |
| Gro/TLE/Grg | (3) |
| AP-1 (Jun/Fos) | (4) |
| CREB1 | (5) |
