## Supplementary material for "Identification of mammalian transcription factors that bind to inaccessible chromatin": Table S7

**Table S6.** Summary of predicted/validated pioneer factors in humans in the literature.

| Cell line | Type | Source | Identifier | Reference |
| --- | --- | --- | --- | --- |
| CH12.LX | DNase | ENCODE | ENCFF001PGJ, ENCFF001PGR, ENCFF001PGO, ENCFF001PGT, ENCFF001PGN, ENCFF001PGL, ENCFF001PGM, ENCFF001PGK, ENCFF001PGQ, ENCFF001PGP, ENCFF001PGS | (1) |
| G1E | DNase | ENCODE | ENCFF001LUD, ENCFF001LUH | (1) |
| MEL | DNase | ENCODE | ENCFF001OMV, ENCFF001OMW, ENCFF001PUL, ENCFF001PUM, ENCFF001PUJ, ENCFF578DTE, ENCFF158XRX | (1) |
| ES-E14 | DNase | ENCODE | ENCFF001PIR, ENCFF001PIW, ENCFF001PIT, ENCFF001PIS, ENCFF001PIV , ENCFF001PIU, ENCFF001PIX | (1) |
| C2C12 | ATAC-seq | GEO | GSM1972411 | (2) |
| G1E-ER4 | DNase | GEO | GSM746567 | (3) |
| E14TG2a.4 | ATAC-seq | GEO | GSM4433145 | (4) |

1. Consortium,T.E.P. (2012) An integrated encyclopedia of DNA elements in the human genome. *Nature*, **489**, 57–74.

2. Dell’Orso,S., Wang,A.H., Shih,H.Y., Saso,K., Berghella,L., Gutierrez-Cruz,G., Ladurner,A.G., O’Shea,J.J., Sartorelli,V. and Zare,H. (2016) The Histone Variant MacroH2A1.2 Is Necessary for the Activation of Muscle Enhancers and Recruitment of the Transcription Factor Pbx1. *Cell Rep.*, **14**, 1156–1168.

3. Wu,W., Cheng,Y., Keller,C.A., Ernst,J., Kumar,S.A., Mishra,T., Morrissey,C., Dorman,C.M., Chen,K.B., Drautz,D., *et al.* (2011) Dynamics of the epigenetic landscape during erythroid differentiation after GATA1 restoration. *Genome Res.*, **21**, 1659–1671.

4. Djeghloul,D., Patel,B., Kramer,H., Dimond,A., Whilding,C., Brown,K., Kohler,A.C., Feytout,A., Veland,N., Elliott,J., *et al.* (2020) Identifying proteins bound to native mitotic ESC chromosomes reveals chromatin repressors are important for compaction. *Nat. Commun. 2020 111*, **11**, 1–15.
